# Healing Tissues From the Inside Out: Infusible Biomaterial for Targeting and Treating Inflammatory Tissues via Intravascular Administration

**DOI:** 10.1101/2020.04.10.028076

**Authors:** Martin T. Spang, Ryan Middleton, Miranda Diaz, Raymond Wang, Jervaughn Hunter, Joshua Mesfin, Tori S. Lazerson, Saumya Bhatia, James Corbitt, Gavin D’Elia, Gerardo Sandoval-Gomez, Rebecca Kandell, Takayuki Kato, Sachiyo Igata, Colin Luo, Kent G. Osborn, Pedro Cabrales, Ester Kwon, Francisco Contijoch, Ryan R. Reeves, Anthony N. DeMaria, Karen L. Christman

## Abstract

Biomaterials, such as extracellular matrix (ECM) hydrogels, have been widely used in preclinical studies as injectable tissue engineering therapies; however, injectable therapies are limited as they can cause localized trauma or organ perforation. We have developed a new ECM therapy, the low molecular weight fraction derived from decellularized, digested ECM, for intravascular infusion. This new form of ECM can be infused after injury, specifically localize to injured tissues by coating the leaky microvasculature, and promote cell survival and tissue repair. In this study, we show the feasibility and targeting of intravascular ECM infusions using models of acute myocardial infarction (MI), traumatic brain injury, and pulmonary arterial hypertension. Furthermore, safety and efficacy were demonstrated in small and large animal models of acute MI following intracoronary infusion, which included using a clinically-relevant catheter in the large animal model. Functional improvements, specifically reduced left ventricular volumes and improved wall motion scores were observed after ECM infusions post-MI. Genes related to tissue repair and inflammation were differential expressed in response to ECM infusions. This study shows proof-of-concept for a new paradigm of delivering pro-healing ECM biomaterials via intravascular infusion to heal tissue from the inside out.

---

Extracellular matrices (ECM) derived from decellularized tissues have shown promising results as tissue engineering scaffolds as an acellular strategy for regenerative medicine^1–5^. In particular, decellularized ECM can be processed via enzymatic digestion into inducible hydrogels that can be injected for minimally-invasive delivery to tissues^6–9^. These degradable hydrogels are pro-survival, immunomodulatory, and promote neovascularization, among a host of other pro-regenerative functions^2,10–12^.

Current decellularized ECM biomaterials are limited to surgically implanted patches or localized injections. We aimed to develop a new form of ECM that could be delivered via jntravascular infusion to target leaky vasculature, allowing for less invasive, more evenly distributed, and potentially earlier delivery since tissue access is not a concern. Acute delivery following injury could reduce cell death, promoting tissue salvage and maintenance. This delivery modality is not amenable to current ECM hydrogels because of their physical properties, which would not allow for passage into the gaps of leaky endothelium at the site of injury or disease.

## RESULTS

### Infusible ECM can be isolated and is capable of gelation

A myocardial ECM hydrogel was selected as a starting material (Fig. 1a-d) for its demonstrated efficacy in preclinical models^7-10,13^ and translational proof-of-concept in a Phase I clinical trial^14^. While this material is injectable, it is a translucent suspension (Fig. 1d), which contains large submicron particulate, which would be too large to pass through leaky vasculature at sites of acute injury and/or inflammation^15,16^.

**Figure 1:**
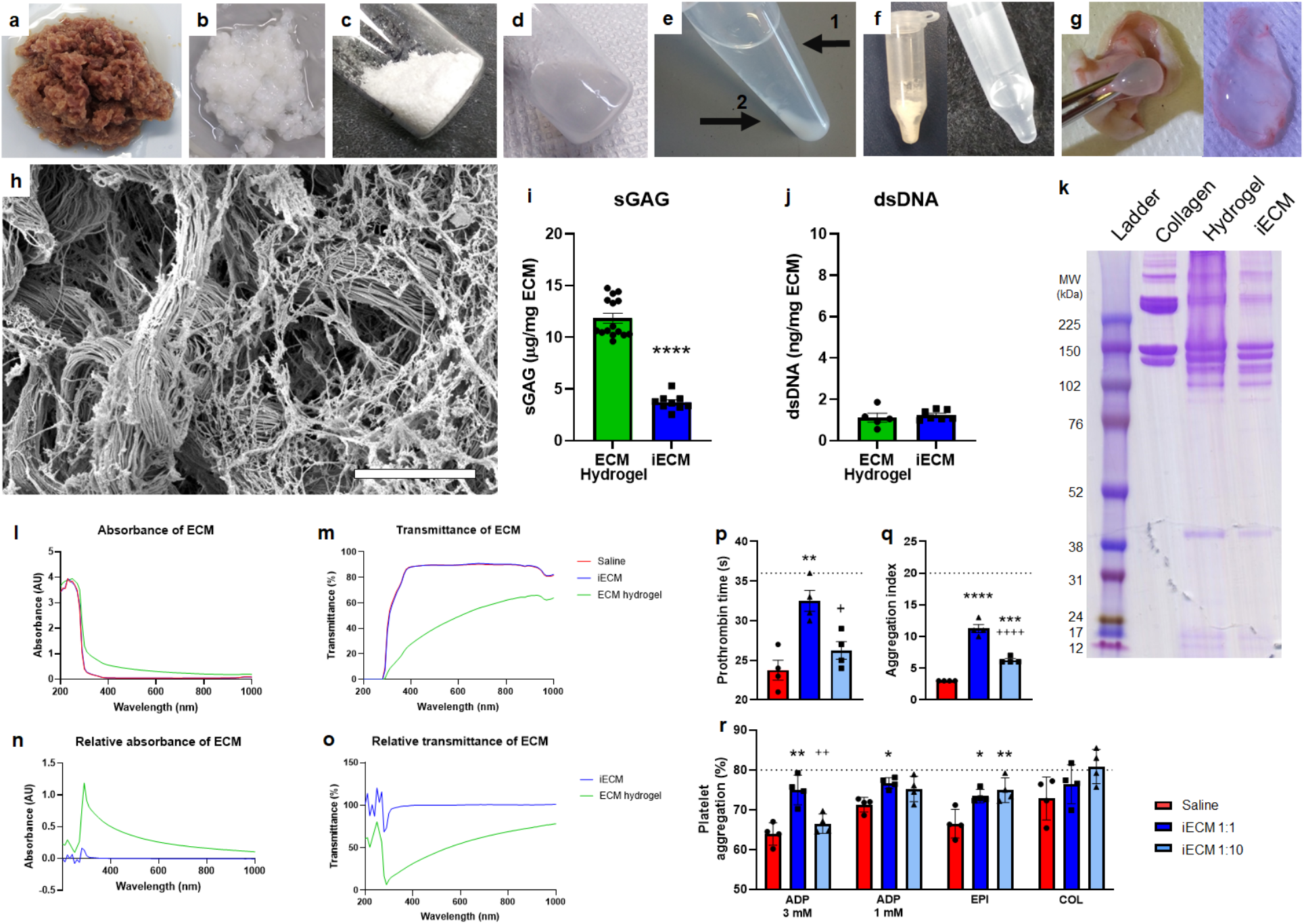
Generation and architecture of infusible extracellular matrix (iECM). **a**, Isolated left ventricular myocardium is cut into pieces. **b**, Decellularized ECM after continuous agitation in 1% sodium dodecyl sulfate. **c**, Lyophilized and milled ECM. **d**, Liquid digested ECM hydrogel. **e**, Fractionated ECM hydrogel after centrifugation; (1) supernatant low molecular weight fraction (iECM) and (2) high molecular weight pellet. **f**, Lyophilized (left) and resuspended (right) iECM. **g**, Subcutaneous injection and gelation of iECM. **h**, Scanning electron microscopy image showing nanofibrous architecture of iECM following gelation. Scale bar is 5 μm. Characterization of iECM, **i**-**o**. **i**, Sulfated glycosaminoglycan (sGAG) content was significantly lower in iECM vs ECM hydrogel. **j**, Double stranded DNA (dsDNA) content was not significantly different between iECM and ECM hydrogel. **k**, Polyacrylamide gel electrophoresis of ladder (Full-Range RPN800E, lane 1), collagen (lane 2), ECM hydrogel (lane 3), and iECM (lane 4), showing depletion of high molecular weight (>200 kDa) peptides/proteins in iECM. **l**-**o**, Optical measurements of iECM vs liquid ECM hydrogel. iECM showed minimal differences in optical properties from saline whereas the ECM hydrogel possessed increased absorbance and decreased transmittance. **l**, Absorbance sweep of iECM, ECM hydrogel, and saline. **m**, Calculated transmittance of iECM, ECM hydrogel, and saline. **n**, Relative absorbance sweep of iECM and ECM hydrogel. **o**, Relative calculated transmittance of iECM and ECM hydrogel. Hemocompatibility of infusible extracellular matrix (iECM) with human blood and platelet rich plasma. **p**, Prothrombin time. **q**, Red blood cell aggregation index. **r**, platelet aggregation following addition of agonists: adenosine diphosphate (ADP), epinephrine (EPI), collagen (COL). Standard ranges for each parameter are indicated between or below dashed lines. * is relative to saline, and + is relative to iECM. Data are mean ± SEM. *p<0.05, **p<0.01, ***p<0.001, ****p<0.0001

To generate an intravascularly infusible ECM, an optimized centrifugation, dialysis, and sterile filtration protocol was developed to remove the high molecular weight peptides/proteins (Fig. 1e). The supernatant was isolated from the high molecular weight pellet and dialyzed, yielding the low molecular weight fraction, infusible ECM (iECM), which could then be sterile filtered, lyophilized for long-term storage, and resuspended for injection/infusion (Fig. 1f). While the iECM did not gel at body temperature *in vitro* like the ECM hydrogel counterpart, it was still capable of gelation in tissue after subcutaneous injection (Fig. 1g). Scanning electron imaging of the gel demonstrated a nanofibrous architecture (Fig. 1h) similar to most ECM hydrogels^17^.

When comparing iECM to the ECM hydrogel, sulfated glycosaminoglycan content was significantly decreased (Fig. 1i), whereas double stranded DNA content was not different (Fig. 1j). Gel electrophoresis showed a depletion of high molecular weight proteins (>200 kDa), which were likely removed during the fractionating process (Fig. 1k). Across most wavelengths tested, the optical properties of iECM were nearly identical to saline, whereas the liquid ECM hydrogel showed increased absorbance and decreased transmittance (Fig. 1l-o), reinforcing the absence of large particles in iECM.

### Infusible ECM is hemocompatible

As iECM was derived from the ECM hydrogel, it was expected that iECM would be hemocompatible. While it is counterintuitive to think that decellularized ECM could be hemocompatible since exposed ECM initiates clotting *in vivo,* we previously demonstrated that the ECM hydrogel was hemocompatible^9^. iECM was tested with human blood using the highest likely scenario of 1:1 iECM to human blood and a physiologically relevant concentration of 1:10 iECM, given the immediate dilution with blood following infusion. Nearly all prothrombin times, red blood cell aggregation indices, as well as platelet aggregation with agonists fall within standard physiological ranges (Fig. 1p-r)^18,19^. Fibrinogen and platelet concentrations were unaffected by the addition of iECM (Extended Data Fig. 1).

### ECM infusions target regions of injury in multiple preclinical models

Three preclinical models were selected to cover diverse conditions in which leaky vasculature is observed, specifically acute myocardial infarction (MI), traumatic brain injury (TBI), and pulmonary arterial hypertension (PAH), which represent both acute and chronic injuries while also covering ischemic, traumatic, and pressure overload injuries. iECM was delivered intravascularly in all models with simulated intracoronary infusion in the MI model and intravenous (IV) infusion in the TBI and PAH models. In the acute MI model (Fiz 2a-h), iECM was localized to the infarcted region (Fig. 2a,b). Minimal material was observed in the border zone (Fig. 2c), and no material was seen in the remote myocardium (Fig. 2d), including the septum and right ventricle. In the TBI model (Fig. 2i-n), material was also localized to the region of trauma (Fig. 2i,j) and no material was observed in remote brain (Fig. 2k). In the PAH model (Fig. 2o-s), material was localized throughout the lungs (Fig. 2o-p), as the PAH model affects the entire organ. All results observed histologically were also confirmed following near-infrared fluorescent scans (Fig. 2g,h, 2n, 2s). Biodistribution analysis suggested excess iECM accumulates in the liver and kidneys (Extended Data Fig. 2). As controls, saline or a small tagged non-gelling peptide (trilysine) control were infused and showed no retention (Fig. 2e,f, 2l, 2q, Extended Data Fig. 3), with the majority going to the kidneys (Extended Data Fig. 2).

**Figure 2:**
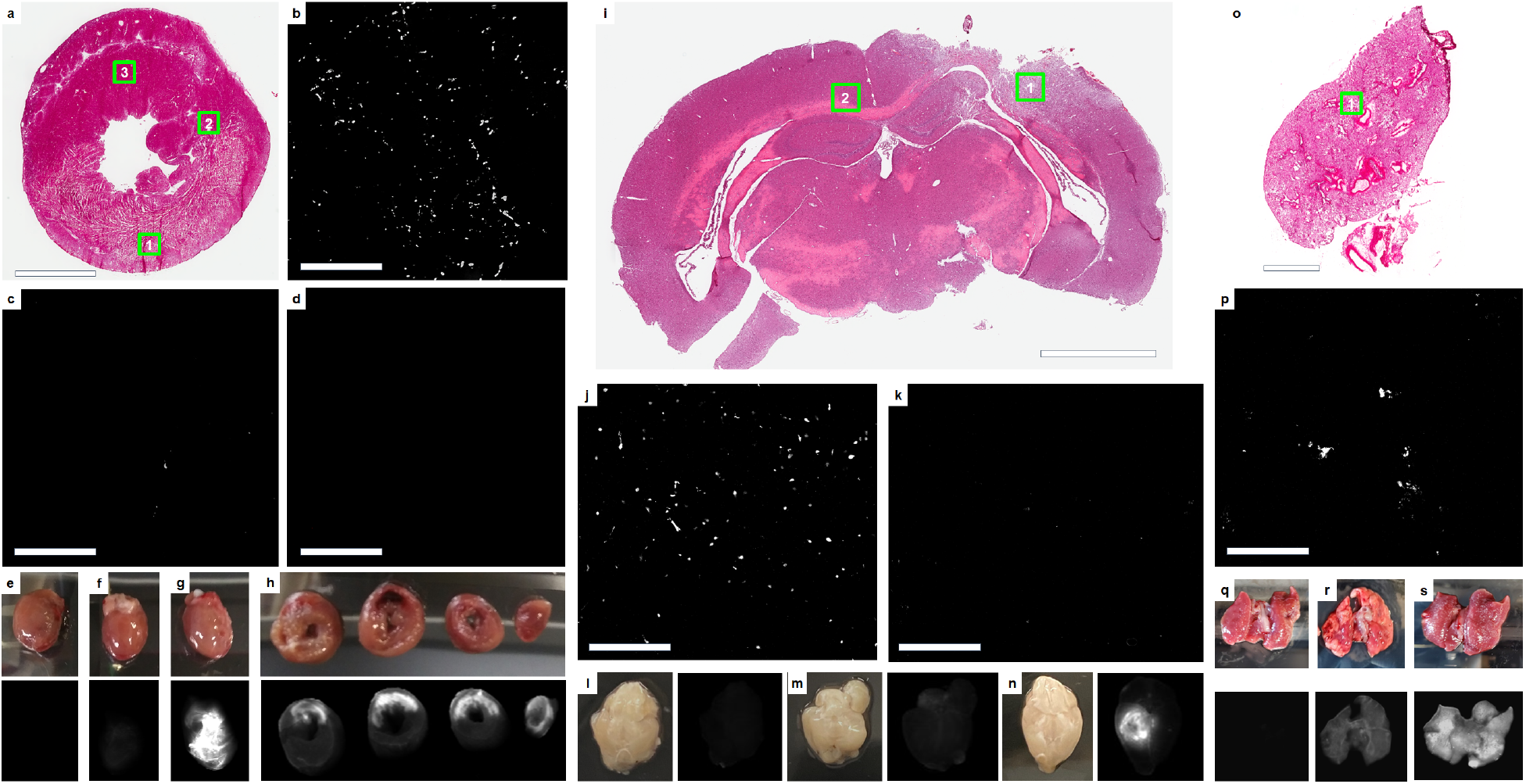
iECM infusions target injured tissues. **a**, H&E short axis section of an acute MI heart following iECM infusion, scale bar 3 mm. **b**-**d**, Fluorescence images for locations shown in insets in **a** of infarcted myocardium (**b**, 1), border zone (**c**, 2), and remote myocardium (**d**, 3), scale bars are 200 μm. **e**-**h**, Heart near-infrared scans of saline without dye (**e**), trilysine conjugated with VT750 (**f**), and iECM conjugated with VT750 (**g**). **h**, Short axis slices of an iECM infused heart from the base to the apex (left to right), infarct regions oriented in the upper portion of the slices. iECM tracking using VivoTag®-S 750 (VT750) 24 hours following infusion and MI procedure. **i,** H&E section of a brain following traumatic brain injury (TBI) and iECM infusion. **j**-**k**, Fluorescence images for locations shown in insets in **i** of injured brain (**j**, 1) or remote brain (**k**, 2). Near-infrared scans of brain infused with saline without dye following TBI (**l**), healthy brain infused with iECM conjugated with VT750 (**m**), and brain infused with iECM conjugated with VT750 following TBI (**n**). **o,** H&E section of a lung following monocrotaline injury, model of pulmonary arterial hypertension (PAH), and iECM infusion. **p**, Fluorescence image for location shown in inset in **o** of injured lung (1). Lung near-infrared scans of lung infused saline without dye following PAH (**q**), healthy lung infused with iECM conjugated with VT750 (**r**), and lung infused with iECM conjugated with VT750 following PAH (**s**).

Across all models, iECM was distributed specifically to injured tissues, suggesting leaky vasculature is required for material retention. To confirm this, iECM infusions were performed in healthy animals without MI (Extended Data Fig. 4), TBI (Fig. 2m), or PAH (Fig. 2r) as well in rats with a chronic MI, which does not have leaky vasculature (Extended Data Fig. 4). All of these animals showed minimal matrix retention in the target tissue; however, material was observed in the kidneys and urine, suggesting almost immediate elimination (Extended Data Fig. 4).

Using the MI model, iECM showed a dose-dependent retention and distribution, increasing from 6 to 10 mg/ml, but with no appreciable increase from 10 to 12 mg/ml (Extended Data Fig. 5). However, due to manufacturing limitations, specifically filtration, 10 mg/ml was the highest concentration that we could routinely produce. Furthermore, iECM was retained for approximately 3 days following infusion (Extended Data Fig. 6).

### Infusible ECM shows co-localization with endothelial cells and reduces vascular leakage

For safety evaluation, we wanted to confirm that iECM was not blocking the lumen of blood vessels. We confirmed that iECM did not block arterioles (Fig. 3a), but we then observed that the iECM aggregates were overlapping with the microvasculature network, specifically endothelial cells (Fig. 3b). Upon further observation, confocal imaging suggested that material was present within the gaps between endothelial cells, resembling a fibrous matrix, and/or coating the inner lining, but not blocking the lumen of capillaries (Fig. 3c-g), leading to the hypothesis that iECM infusions were reducing pathological tissue permeability by filling in the gaps of leaky vasculature. In line with this hypothesis, iECM infusions significantly decreased leakage of tagged BSA into the infarct (Fig. 4h-j), suggesting iECM reduced tissue permeability. However, this did not affect neutrophil emigration (Extended Data Fig. 7).

**Figure 3:**
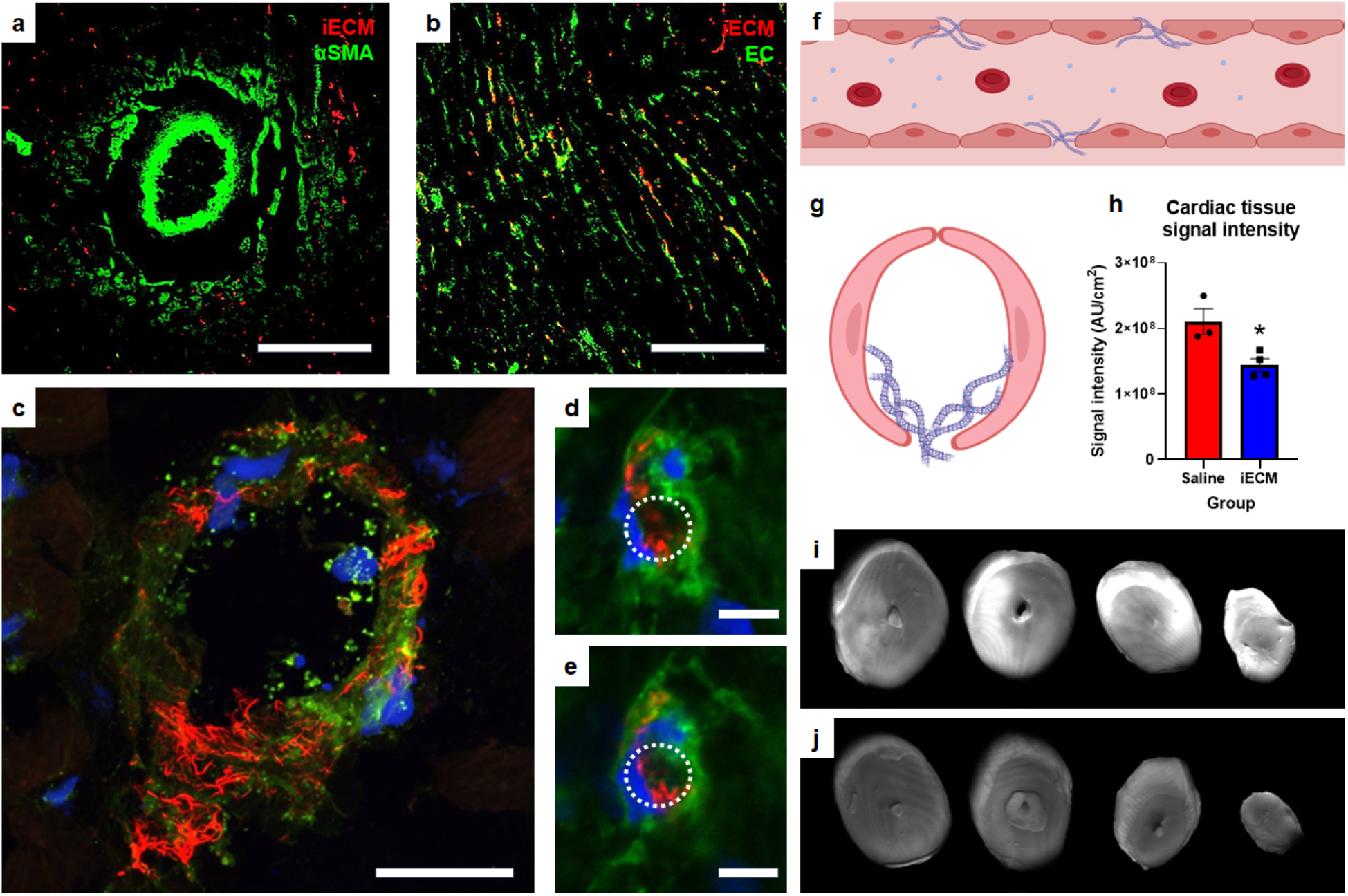
Staining for vasculature and iECM localization post-infusion and MI, showing colocalization of iECM with the microvasculature as opposed to large vessels. iECM was pre-labeled and visualized in red in all parts of Fig. 3. **a**, Staining for arterioles using anti-alpha smooth muscle actin (αSMA) in green. (**b**-**e**), Staining for endothelial cells using isolectin in green. Scale bar is 200 μm for **a**-**b**. **c**, iECM does not block the lumen of an arteriole but fills in the gaps in the endothelium. Scale bar is 25 μm. **d**,**e**, Representative sequential z-stack images of a capillary showing that iECM coats the lumen while not blocking it. Lumen traced with dotted white lines. Scale bar is 5 μm. **f**, Diagram of longitudinal axis view of iECM fibers coating a leaky vessel. **g**, Diagram of short axis view of iECM coating a capillary. **h**-**i,** Fluorescent scans of hearts following MI, intracoronary infusion (ICI) of saline (**h**) or iECM (**i**), and IV infusion of fluorescent bovine serum albumin (IV BSA). **j**, Quantified cardiac tissue signal intensity following MI, ICI, and IV BSA, suggesting iECM infusions decrease tissue permeability. Data are mean ± SEM. *p<0.05.

**Figure 4:**
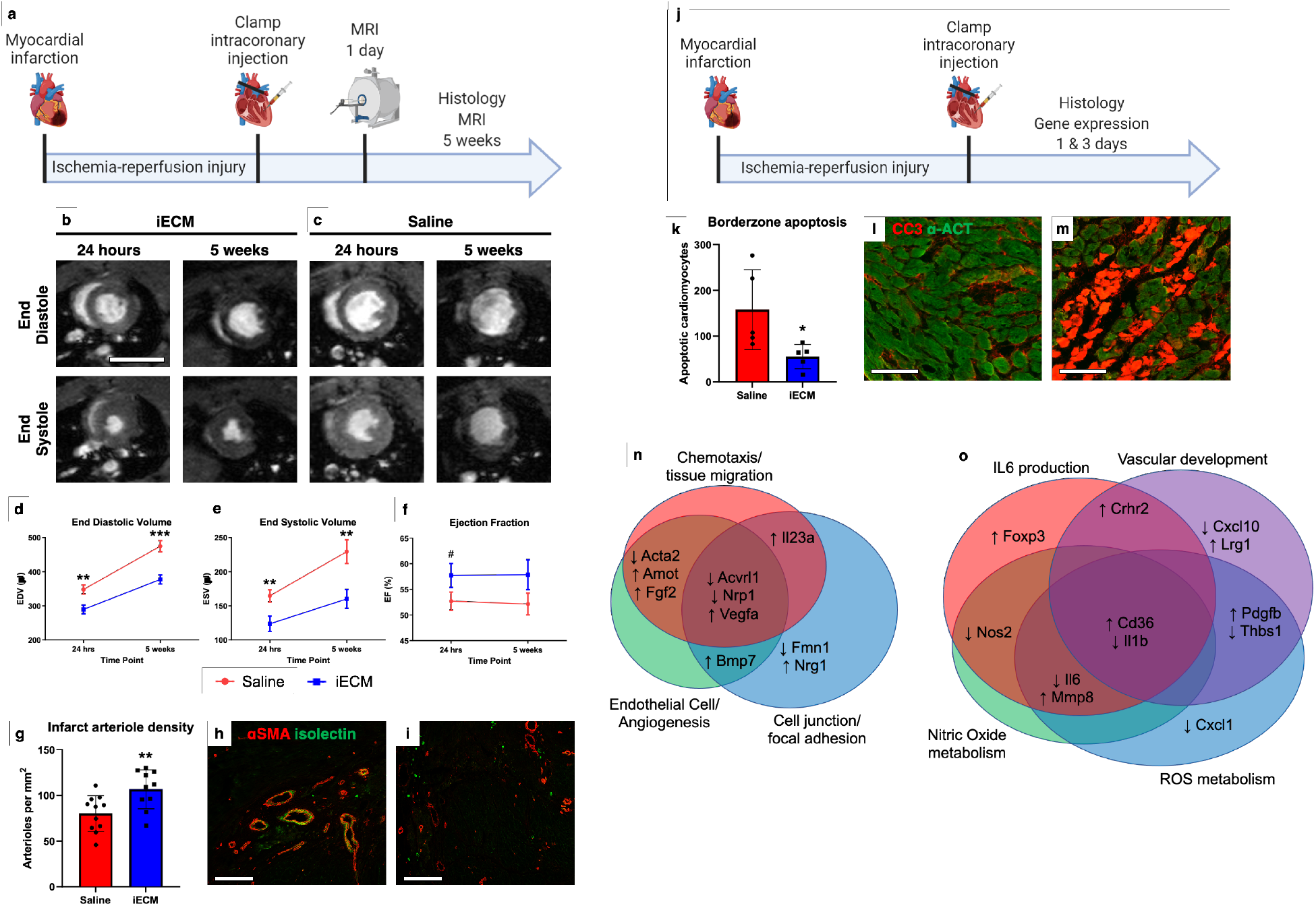
Infusible extracellular matrix (iECM) significantly improved cardiac function post-MI. **a**, Timeline of survival study. **b**-**c**, Representative magnetic resonance images at 24 hours and 5 weeks post-injection of iECM (**b**) or saline (**c**). Scale bar 10 mm. **d**-**f**, iECM infusions preserve left ventricle volumes at 24 hours and 5 weeks post-MI, end diastolic volume (**d**, EDV), end systolic volume (**e**, ESV), ejection fraction (**f**, EF). **g**, Infarct arteriole density increased with iECM infusions at 5 weeks post-infusion, promoting neovascularization. Representative images of iECM (**h**) and saline (**i**) infused hearts, alpha smooth muscle actin (αSMA) in red for arterioles and isolectin in green for endothelial cells. Scale bar 250 μm. **j**, Timeline of acute mechanisms of repair study. Mechanisms of repair were evaluated through histology (**k**-**m**) and gene expression (**n**-**o**) analyses. **k**, Number of cardiomyocytes undergoing apoptosis significantly decreased in the border zone of iECM infused hearts at 3 days post-infusion. Representative images of iECM (**l**) and saline (**m**) infused hearts, cleaved caspase 3 (CC3) for apoptosis in red, and alpha actinin (α-ACT) for cardiomyocytes in green. Scale bar 100 μm. **n**-**o**, Gene expression analysis from NanoString analysis at acute timepoints, suggesting genes related to chemotaxis/tissue migration, endothelial cells/angiogenesis, and cell junction/focal adhesion proteins are differentially expressed 1 day following iECM infusions and genes related to Interleukin 6 (IL6), vascular development, nitric oxide metabolism, and reactive oxygen species (ROS) metabolism are differentially expressed at 3 days following infusion. Data are mean ± SEM.*p<0.05, **p<0.01, ***p<0.001, ^#^p<0.10.

### Infusible ECM mitigates negative left ventricular remodeling in a rat acute MI model

To test the efficacy of iECM in an acute MI model, rats were infused with iECM or saline following occlusion-reperfusion (Fig. 4a-c). At 24 hours following infusions, iECM significantly decreased LV volumes, both end-systolic (ESV, Fig. 4d) and diastolic volumes (EDV, Fig. 4e), compared to the saline control alongside a trending increase in ejection fraction (EF, Fig. 4f). We confirmed that this was not a result of differences in infarct size between groups (Extended Data Fig. 8a-d) suggesting similar initial injuries. This significant decrease in LV volumes of iECM animals was maintained at 5 weeks post-infusion as well.

### Infusible ECM is pro-survival and promotes tissue repair

As ECM hydrogels have previously shown to promote tissue repair, including increases in vascularization, being pro-survival, and modulation of inflammation^10,20^, we hypothesized that iECM would likewise have similar pro-reparative effects. At 5 weeks post-infusion, iECM significantly increased infarct vascularization over controls (Fig. 4g-i). Infarct fibrosis and interstitial fibrosis were not significantly different between groups (Supplemental Fig. 8e-g). At 3 days post-infusion (Fig. 4j), iECM significantly decreased the number of cardiomyocytes undergoing apoptosis in the border zone region (Fig. 4k-m). We observed significant differential expression of 11 and 23 genes at 1 day and 3 days post-infusion, respectively. Gene Ontology (GO) pathway enrichment analysis suggested pathways, including angiogenesis, cell-substrate, reactive oxygen species (ROS) and nitric oxide (NO) metabolism, and interleukin 6 (IL6) and other cytokine signaling as illustrated by the Venn diagram of differentially expressed genes in Figure 4n,o. The pathway distribution and full gene list are in Extended Data Tables 1 & 2. Heat maps of differentially expressed genes and the top 30 pathways from GO analysis can be viewed in Extended Data Figure 9 and Extended Data Tables 3 and 4, respectively.

### Infusible ECM is amenable to intracoronary infusion with a clinically-relevant catheter

Given the promising results in the small animal acute MI model, we tested if iECM could also be delivered using a clinically-relevant infusion catheter in a large animal model. To ensure that iECM would pass through a catheter, pre-gel complex viscosity was measured (Fig. 5a). The pre-gel complex viscosity of iECM was an order of magnitude lower than the liquid ECM hydrogel, but still higher than saline (Fig. 5a). Despite the increased viscosity, iECM was able to pass through a catheter with an internal diameter of 0.36 mm. To assess clinical feasibility, a balloon infusion catheter was used to induce MI and infuse tagged iECM (Fig. 5b). As seen in the small animal model, iECM was observed in the infarct region in the porcine MI model (Fig 5c,d). Likewise, iECM was distributed throughout the infarcted myocardium and was not observed in the border zone or remote myocardium (Extended Data Fig. 10). Satellite organs did not show any signs of material, or acute ischemia or inflammation (Extended Data Fig. 11, Extended Data Table 5). iECM was observed to similarly line endothelial cells of the infarcted myocardium in the pig model (Fig. 5e,f). No signal was observed in the infarct region with a small peptide control (Extended Data Fig. 10).

**Figure 5:**
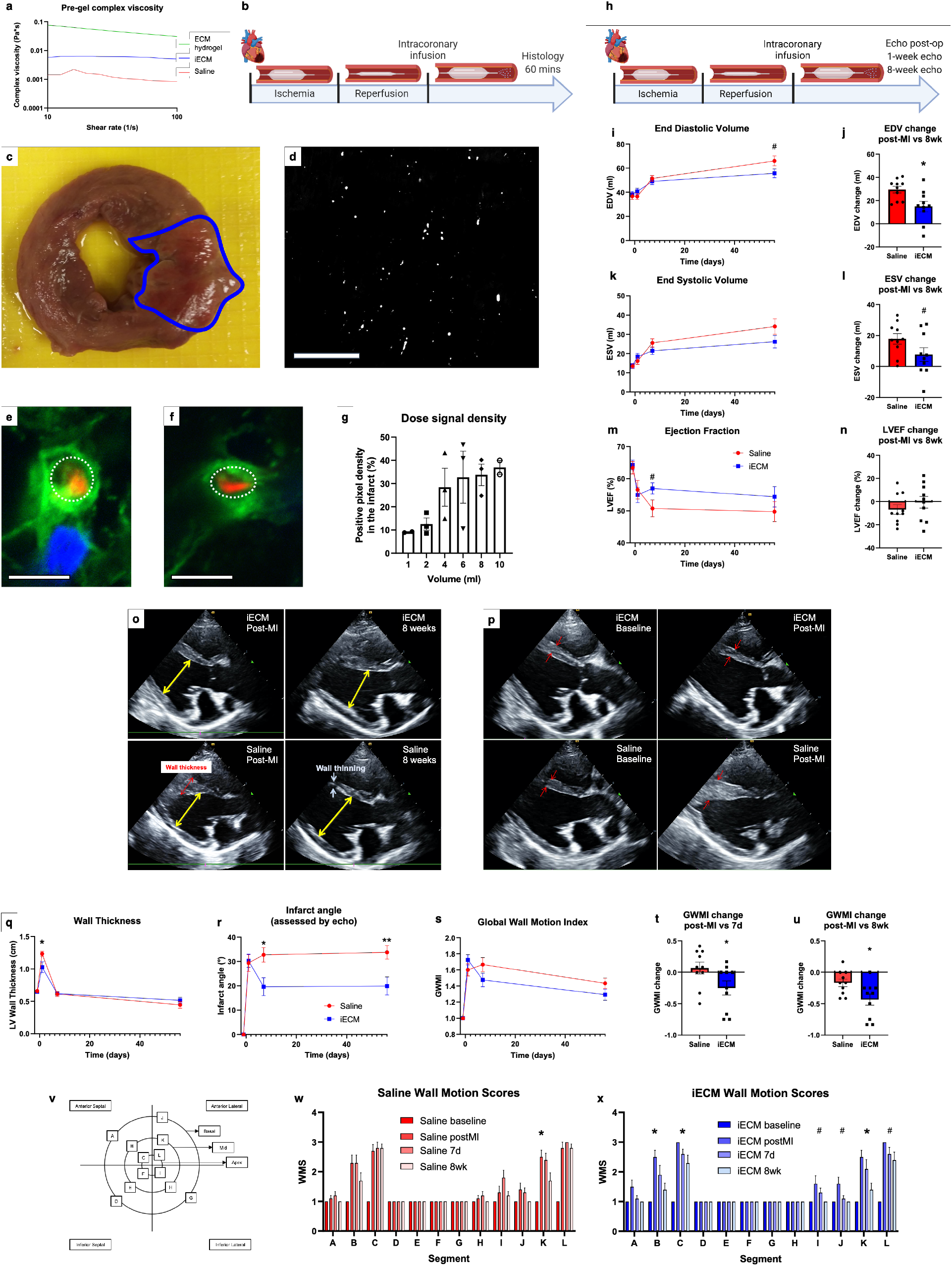
iECM infusions are amenable to intracoronary infusion with a balloon infusion catheter and mitigate negative left ventricular remodeling in a pig acute MI model. **a**, iECM pre-gel complex viscosity is lower than its full ECM hydrogel counterpart, suggesting potential for catheter delivery. **b**, Timeline of catheter feasibility study. **c**, Macroscopic short-axis view of a pig heart post-MI and iECM infusion. Infarct outlined in blue. **d**, Fluorescent image of iECM distribution and retention in a histological section from the same heart. Scale bar 200 μm. **e**,**f**, Representative confocal images of iECM (red) and endothelial cells (green) from iECM infused pigs. iECM lines the endothelial cells of infarcted myocardium, as similarly observed in the rat MI model. Lumen traced with dotted white lines; Scale bars 5 μm. **g**, iECM infusions showed a plateau effect when increasing infusion volumes, as suggested by quantified signal density in the infarct. **h**, Timeline of survival study following induced MI and iECM or saline infusion. End diastolic volume (EDV), end systolic volume (ESV), ejection fraction (EF), left ventricular (LV) wall thickness, infarct angle, global wall motion index (GWMI), and wall motion scores (WMS) were measured before MI (baseline), post-MI, 7 days, and 8 weeks post-MI. **h**-**u**, Changes in EDV, ESV, and EF over time (**i,k,m**) and relative to post-MI (**j,l,n**), suggest iECM infusions mitigate negative left ventricular remodeling. Representatives echocardiography images at end diastole show how iECM infusions mitigate negative LV remodeling from post-MI to 8 weeks post-MI (**o**) and mitigate acute increases in wall thickness post-MI (**p**). Red arrows indicate wall thickness, yellow arrows are used for LV end diastolic measurement. Changes in MI wall thickness (**q**), infarct angle (**r**), GWMI (**s-u**), and WMS segments (**v-x**) over time, showing how iECM infusions preserve LV wall thickness and motion and mitigate infarct expansion. Wall motion score diagram used for scoring wall motion (**v**). Data are mean ± SEM. *p<0.05, ^#^p<0.10.

### Infusible ECM mitigates negative left ventricular remodeling in a porcine acute MI model

To optimize material retention, volumes ranging from 1 ml to 10 ml were infused following MI, demonstrating a plateau effect at 4 ml, suggesting that any additional infused iECM would not be retained, as the microvasculature may be saturated (Fig. 5g). Subsequently, pigs underwent MI, iECM or saline infusion, and serial echocardiography (echo) (Fig. 5h-x, Extended Data Fig. 12). Immediately post-MI, EDV, ESV, EF, and infarct angle, a measure of infarct area (Extended Data Fig. 12h), were not significantly different between groups, suggesting a similar injury between groups. EDV, ESV, and EF were measured over time (Fig. 5i,k,m) showing iECM significantly reduced the expansion in EDV from immediately post-MI to 8 weeks later compared to saline (Fig. 5j,o) with a similar trending improvement in the change in ESV (5l) and significant improvement in the change in LV systolic dimension from post-MI to 8 weeks post-MI (Extended Data Fig. 12e). While the change in EF was not significantly different (Fig. 5n), there was significant improvement in the change in fractional shortening (Extended Data Fig. 12g).

Shortly post-MI and iECM infusion, LV wall thickness was significantly decreased (Fig. 5p,q), while infarct angle was significantly reduced at 7 days and 8 weeks post-MI (Fig. 5r) compared to saline. Global wall motion index scores also showed similar improvements (Fig. 5s) with significant differences in the change from post-MI to 7 days (Fig. 5t) and 8 weeks (Fig. 5u). This was attributed to significant improvements in infarcted wall segments (B, C, K, and L) over time, whereas no improvements were observed for saline infused hearts (Fig. 5v-x). ECG and Holter monitoring analysis showed no differences in arrythmias between groups.

## Discussion

In this study, we developed a new form of decellularized ECM for intravascular infusion as a new acellular strategy for regenerative medicine. We demonstrated feasibility of delivering iECM intravascularly to target injured and diseased tissue with leaky vasculature, showing broad applicability in acute MI, TBI, and PAH models. iECM was hemocompatible and we found no evidence of embolization, demonstrating the material can be safely delivered intravascularly. Furthermore, in small and large animal acute MI models, therapeutic efficacy was observed, indicating the potential for clinical translation. The material was only observed in injured tissues with leaky vasculature, demonstrating that iECM retention and therapeutic efficacy is conserved across species and injury models.

Our initial hypothesis was that iECM would pass through the gaps in the endothelial cell layer in the leaky vasculature and into the injured tissue as was previously seen with an infusible alginate material^21^. Instead, we found that iECM coats and/or fills in the gaps between endothelial cells (Fig. 3), specifically in regions of damaged tissues, and therefore represents a new type of targeted delivery for pro-healing biomaterials. This localization led to a reduction in vascular permeability (Fig. 3h-j), with potential reduction in tissue edema, which, in the case of MI, is known to contribute to cardiac cell dysfunction and death^22,23^. As observed in the pig MI model, wall thickness was significantly reduced post-MI, suggesting a barrier to fluid and/or small proteins. As myocardial edema begins during ischemia and quickly expands following reperfusion^24^, acute iECM infusions could prevent tissue edema and further tissue damage from reperfusion injury. Tissue edema is also observed in numerous pathologies with leaky vasculature, including TBI^25,26^ and PAH^27,28^, suggesting iECM could be a novel therapeutic approach for such conditions.

To demonstrate the therapeutic of potential of iECM we extensively tested it in both small and large animal MI models, including the use of clinically relevant catheter procedures in the latter. iECM significantly reduced both end-diastolic and end-systolic volumes one day after MI and infusion in rats. This difference was maintained, but did not further improve out to 5 weeks post-infusion (Fig 4d,e), essentially creating a better baseline. This is in line with the observed degradation time of iECM, approximately 3 days, where there is an initial therapeutic effect. Therapeutic efficacy was also observed in the porcine model where iECM mitigated an increase in EDV from post-MI to 8 weeks post-MI, reinforcing that iECM infusions reduce negative LV remodeling. Significant improvements in infarct angle and wall motion were also observed in the porcine model, further suggesting localized therapeutic efficacy and improvements in regional function.

In terms of biological activity, ECM hydrogels have been shown to create a pro-remodeling versus a pro-inflammatory environment, leading to increases in vascularization and cell survival^7,10,31^. We found decreased numbers of apoptotic cardiomyocytes and increased arteriole density with the iECM infused rat hearts. However, based on the finding that iECM coats endothelial cells as opposed to entering the tissue, the mechanisms leading to improved survival and vascularization are likely different between the iECM and ECM hydrogels. From gene expression analyses, angiogenic, endothelial cells, and cell migration/recruitment pathways were expected and dominated the pathway analysis 1 day following iECM infusions. It was unexpected to see several pathways around focal adhesions and cell-substrate junctions. The focal adhesion and cell-substrate pathways may suggest how the treated endothelial cells adapt in response to iECM infusions, as endothelial cells would be directly exposed immediately post-infusion. At 3 days post-infusion, ROS and NO metabolism and IL6 signaling emerged. It is possible that iECM infusions encourage healthy ROS metabolism or the iECM itself provides an ROS sink and/or protection for the surrounding cells as has been shown with the ECM hydrogel^32^, leading to decreased apoptosis. IL6 downregulation at 3 days post-MI would suggest that iECM infusions encourage this pro-regenerative switch to happen earlier, reducing negative LV remodeling^33^.

Even though iECM is derived from an ECM hydrogel, it could be more appropriate to compare iECM to peptide therapeutics given its delivery modality. This comparison to peptides is evident in the clearance of non-retained iECM, which like other peptide therapeutics^29,30^ appears to be rapidly excreted by the kidneys (Extended Data Fig. 2). However, iECM has advantages over traditional peptide therapeutics due to increased retention through potential binding and gelation in the gaps of leaky vasculature (Fig. 3c). This could shield the material from proteolysis, allowing for increased tissue retention and therapeutic effects over peptides therapeutics^30^.

ECM infusions are a versatile new platform for the regenerative medicine and tissue engineering field. In this study, we chose to focus on the cardiac application of ECM infusions; however, infusible ECM could be generated from any type of decellularized tissue, and ECM infusions could be applied to many diseases or injuries where there is endothelial cell injury or dysfunction. Whereas traditional tissue engineering approaches have focused on scaffolds and/or cells to replace or repair damaged tissue, infusible ECM therapies could shift the focus towards treating the microvasculature and healing tissues from the inside out to improve tissue function.

## Acknowledgements

Funding was provided by the NIH NHLBI (R01HL113468, R43HL150917). MTS was supported by NIH NHLBI T32HL105373 and an AHA pre-doctoral fellowship. The authors would like to thank Jesse Placone for his confocal imaging assistance, Pamela Duran for her helpful comments during the manuscript editing process, and the University of California, San Diego Institutional Animal Care and Use Program veterinary staff for their assistance with large animal procedures and safety. Schematic figures were created with BioRender.com.

## Author contributions

KLC obtained the funding. MTS, KLC, RRR, and AND conceptually designed the studies. MTS, KLC, and AND interpreted the results. CL performed small animal surgeries. SI performed large animal echocardiography. RRR performed large animal surgeries. KGO performed large animal necropsy and histopathology analysis. MTS, RM, MD, RW, JH, JM, TSL, SB, JC, RK, and GS processed tissue samples and performed analyses. RM, RW, JH, JM, TK, and CL assisted with large animal surgeries. MD and RW assisted with gene expression analyses. PC designed and performed hemocompatibility analysis. EK and FC provided consultation, experimental design, and data evaluation related to imaging. MTS and KLC drafted the manuscript. All authors edited the manuscript.

## Additional information

Supplementary information is available for this paper online.

Correspondence and requests for materials should be addressed to KLC.

