## Supplementary Materials for "Healing Tissues From the Inside Out: Infusible Biomaterial for Targeting and Treating Inflammatory Tissues via Intravascular Administration"

### METHODS

#### Materials

Unless specified, all materials were purchased from Sigma Aldrich.

#### Material production

Extracellular matrix (ECM) hydrogel was generated based on previously described protocols[1-3]. In brief, fresh hearts were harvested from adult Yorkshire farm pigs (30-45 kg), and the left ventricular (LV) myocardium was isolated. Major vessels and connective tissue were removed, and the remaining tissue was minced. Tissue was decellularized in 1% sodium dodecyl sulfate (SDS) in phosphate buffer saline (PBS) for 4-5 days, followed by 24 hours of water rinsing to remove detergent. The material was then lyophilized and milled into a fine powder, and subsequently partially enzymatically pepsin digested in 0.1 M hydrochloric acid at 10 mg/ml ECM powder and 1 mg/ml pepsin for at least 48 hours. The material was then neutralized with sodium hydroxide and buffered to match *in vivo* conditions, yielding liquid ECM hydrogel, capable of thermally induced gelation. ECM hydrogel was diluted to 6 mg/ml using PBS.

Next, the liquid ECM hydrogel was centrifuged at 15,000 RCF at 4°C for 45 minutes to separate the high and low molecular weight fractions. The supernatant (infusible ECM, iECM) was isolated from the high molecular weight pellet. The pellet was rinsed with PBS and centrifuged again to increase yield of iECM. To adjust the concentration and ratio of salts, iECM was dialyzed over 48 hours at 4°C in 0.5xPBS, 0.25xPBS, and then twice in deionized water (each solution was replaced approximately every 12 hours), and lyophilized. The iECM was then resuspended at a high concentration (16 mg/mL), passed through 0.22 µm syringe filters (Millipore) into sterile test tubes, lyophilized, weighed, and stored at -80°C for future use. The iECM was then resuspended to appropriate concentration in sterile water at least 30 minutes before injection while on ice.

**Characterization assays**

Characterization of iECM and its full ECM hydrogel counterpart was performed as previously described[2]. Protein size and distribution of iECM and ECM hydrogel were visualized against rat tail collagen I through SDS polyacrylamide gel electrophoresis (PAGE) NuPage Kit (Invitrogen). A 4-12% Bis-Tris gel (Invitrogen) and Full-Range RPN800E ladder (MilliPore Sigma) were used for PAGE. Protein bands were visualized using Imperial Protein Stain. For double stranded DNA (dsDNA) quantification, iECM and ECM hydrogel were digested using Proteinase-K, and DNA was isolated using a NucleoSpin kit (Macherey-Nagel). dsDNA was quantified using a PicoGreen fluorescent reporter (Life Technologies). Sulfated glycosaminoglycans (sGAGs) were quantified using a 1,9-dimethylmethylene blue (DMMB) assay[2] and compared against chondroitin sulfate standards. All values are reported relative to the dry mass of iECM or ECM hydrogel. Optical properties (absorbance) were measured via an absorbance scan on a Spark® microplate reader (Tecan) with 10 nm steps using 1:50 dilutions of iECM and ECM hydrogel (final concentrations of 0.2 mg/ml and 0.12 mg/ml, respectively) alongside saline as a control. Transmittance was calculated from absorbance using the equation, Transmittance = 10^(2-Absorbance). Relative absorbance and transmittance were calculated by subtracting the saline signal values from iECM and ECM hydrogel values.

**Gelation and scanning electron microscopy**

To test gelation, 500 µL of iECM was resuspended to 10 mg/ml and subcutaneously injected in two locations on the back of a female Sprague-Dawley rat. Subcutaneous gels were harvested 5 minutes following injections, dissected from tissue, fixed in 4% paraformaldehyde and 4% glutaraldehyde, dehydrated in ethanol, transferred to isopropyl alcohol, critical point dried using an AutoSamdri 815A automated critical point drier (Tousimis, Rockville, MD), sputter coated (Leica SCD500, Leica, Vienna) with approximately 7 nm of iridium, and imaged at 10,000X under an FEI Quanta 250 scanning electron microscope (Thermo Fisher, Waltham, MA) at 3kV using the in-lens SE1 detector.

**Hemocompatibility**

The interaction between iECM and human blood samples (N=4) was assessed at different concentrations of iECM to whole human blood or platelet rich plasma. A ratio of 1:1 (10 mg/ml iECM) represents the highest possible ratio between blood and iECM, whereas 1:10 (1 mg/ml) represents a more physiologically relevant dilution based on the volume of the coronary vasculature and relevant infusion rate (1 ml/min). Hemocompatibility was assessed as previously described[4-6]. Red blood cell aggregation was measured using 20 µl of blood with a photometric rheoscope (Myrenne Aggregometer, Myrenne Gmbh, Roetgen, Germany) within 4 hours of sampling after adjusting hematocrit to 45% with autologous plasma. The aggregometer measures changes in light transmission following red blood cell shearing, resulting in a decrease in optical signal from red blood cell aggregation. The aggregation time is the reciprocal of the slope (calculated between 0.5 and 2 s after shear has stopped). The aggregation index is the relative surface area above the curve calculated over the first 5 s. Platelet aggregation was measured with isolated platelet rich plasma on a lumi-aggregonometer (Chrono-log). Using the same dilutions as above for sample to platelet rich plasma, high concentration coagulation cascade agonists, including adenosine diphosphate

(ADP), epinephrine (EPI), and collagen (COL) were added (1:200-1:1000 dilutions), and platelet aggregation was measured via absorbance (600-620 nm).

**Animal surgeries**

All procedures in this study were performed in accordance with the guidelines established by the Committee on Animal Research at the University of California, San Diego and the Association for the Assessment and Accreditation of Laboratory Animal Care. All surgeries were performed using aseptic conditions.

**Small animal surgeries**

To induce tissue injury, a rat ischemia-reperfusion model of myocardial infarction (MI)[7,8], a mouse controlled cortical impact (CCI) model of traumatic brain injury (TBI)[9-12], and a rat monocrotaline (MCT) pulmonary arterial hypertension (PAH) model[13,14] were used as previously described. Adult female Sprague-Dawley rats (225-250 g) or C57 black 6 mice aged 8-10 weeks were anesthetized using isoflurane.

To allow for visualization, iECM or trilysine was conjugated with Alexa Fluor™ 568 N-hydroxysuccinimidyl ester to bind free amines (AF568, Invitrogen), as previously described[15]. AF568 was resuspended at 10 mg/ml in dimethyl sulfoxide, and AF568 was added to iECM or trilysine at a 1:100 volume dilution. For biodistribution, iECM and trilysine were conjugated with VivoTag®-S 750 Fluorochrome (VT750, PerkinElmer), similarly to AF568, as previously described[16]. VT750 was resuspended at 10 mg/ml in dimethyl sulfoxide, and VT750 was added to iECM or trilysine, at a 1:100 volume dilution.

To induce acute MI, the left main artery was accessed via a left thoracotomy and was ligated for 35 minutes with a suture. To simulate reperfusion, the suture was released to restore blood flow. To simulate intracoronary (IC) infusion, the aorta was clamped, and iECM, trilysine, or saline was injected into the LV lumen with a 30G needle, forcing the material into the coronary arteries which feed both the left and right side of the heart[17,18]. Infusions were performed within 10 minutes of reperfusion. To determine the effects of dose on retention, hearts were infused with 200 µl of 6, 10, or 12 mg/ml iECM and were harvested at 30 minutes post-infusion (n=2 rats per dose). To determine degradation, rats underwent MI and IC infusion of 200 µl of 10 mg/ml iECM+AF568, were harvested at 30 minutes, 1, 6, 12, and 24 hours and 2, 3, 4, 5, and 7 days post-infusion (n=2 rats per time point). Harvested hearts were rinsed with saline, embedded in Tissue Tek optimal cutting temperature (OCT) compound, and frozen for cryosectioning.

To determine if MI was necessary for iECM retention, IC infusions of 10 mg/ml iECM conjugated with AF568 were performed in healthy rats (no MI), and hearts, kidneys, lungs, liver, kidneys, and spleens were harvested after 1 hour post-infusion and assessed for material (n=3 rats). To assess if retention is possible in a chronic MI, rats underwent MI procedure four weeks prior to infusion. After four weeks post-MI, IC infusions were administered, consisting of either 10 mg/ml iECM conjugated with AF568 (n=2 rats), trilysine conjugated with AF568 (n=1 rat), or saline (n=1 rat). Hearts were harvested 1 hour post-infusion.

To determine MI biodistribution, rats underwent MI and IC infusion of 200 µl of 10 mg/ml iECM+VT750, trilysine+VT750, or saline without VT750 (n=2 rats per group). At 24 hours post-infusion, rats were perfused with saline to reduce background signal, and heart, lung, liver, brain, kidney, and spleen were harvested, rinsed with PBS, and stored at 4°C. Tissues were imaged within 24 hours using an Odyssey imaging system (Li-COR Biosciences).

To induce TBI, a midline incision was made to expose the skull, and a 4 mm diameter craniotomy performed over the right hemisphere between bregma and lambda. The CCI was applied to the exposed dura of the cortex with the ImpactOne (Leica Biosystems) fitted with a stainless steel 2 mm diameter probe at a velocity of 3 m/s and a 2 mm depth. The bone flap was replaced, and the scalp was sutured. To induce PAH, a single subcutaneous/intraperitoneal injection of MCT (60 mg/kg) was delivered. To confirm injury, rats were monitored weekly using echocardiography for changes in Tricuspid annular plane systolic excursion (TAPSE). To determine feasibility in TBI and PAH models, iECM was fluorescently tagged and infused via tail vein injection, and brains or lungs were harvested 1 hour post-infusion. For TBI, iECM (n=3) or saline (n=2) was infused 4 hours following TBI and imaged. Healthy brains (n=2) were also infused via tail vein infusion with iECM and imaged. For PAH, iECM (n=2) or saline (n=2) was infused 5 weeks following MCT injury. Healthy lungs (n=2) were also infused via tail vein infusion and imaged.

To assess vascular permeability, rats underwent MI and IC infusion of 200 µl of saline (n=3) or iECM (n=4). After 30 minutes, rats underwent tail vein injection of Albumin from Bovine Serum (BSA), Alexa Fluor™ 680 conjugate (ThermoFisher Scientific, A34787). After an additional 30 minutes, hearts were harvested and imaged using an Odyssey Odyssey imaging system (Li-COR Biosciences).

To assess efficacy in the acute MI model, rats underwent MI and IC infusion of 200 µl of saline or iECM and then magnetic resonance imaging (MRI) 24 hours and 5 weeks post-infusion (iECM n=10 rats, saline n=11 rats). Animals were arbitrarily assigned with saline and iECM injections being performed each day of surgery. Hearts were harvested within 24 hours of the last imaging time point. Cardiac cine MRI were acquired using an 11.7T Bruker MRI System by Molecular Imaging Inc at the Sanford Consortium for Regenerative Medicine. Rats were anesthetized using isoflurane in oxygen during imaging. Respiratory and electrocardiogram-gated, cine sequences were acquired over contiguous heart axial slices. The following parameters were used: repetition time = 20 ms, echo time = 1.18 ms, flip angle = 30°, field of view = 40 $mm^2$, data matrix size = 200 x 200. Eight or nine 1.5 mm, axial image slices were acquired with a total of 20 cine frames per image slice. ImageJ (NIH) was used to outline the endocardial surface at end diastole and end systole for each slice, defined as the minimum and maximum LV lumen area, respectively. Simpson's method was used to calculate the end-diastolic volume (EDV) and end systolic volume (ESV). Ejection fraction (EF) was calculated as [(EDV-ESV)/EDV] x 100. Investigators were blinded to the groups during image acquisition and analyses.

For gene expression analyses, rats underwent MI and IC infusion of 200 µl of saline or iECM, and hearts were harvested at 1 and 3 days post-infusion (saline 1 day n=5 rats; iECM 1 day, saline 3 day, and iECM 3 day n=6 rats per group). Hearts were sliced into six to seven approximately 1 mm coronal slices using a Rat Heart Slice Matrix (Zivic Instruments). Odd slices were used embedded in OCT for histology analyses, and the even slices were used for gene expression analyses. The LV free wall was isolated from even slices and flash-frozen.

**Large animal surgeries**

Yucatan mini-pigs (35-55 kg) were infarcted and infused using over-the-wire percutaneous transluminal coronary angioplasty (PTCA) dilatation catheters (Boston Scientific), as previously described[19,20]. iECM and trilysine were conjugated AF568 as described above. Pre-anesthesia, pigs were intramuscularly administered 3-5 mg/kg Telazol (Wyesth) or a combination of ketamine-xylazine (4:1, 5 ml). Animals were intubated and maintained under anesthesia using 1-2.5% isoflurane or ketamine-xylazine-atropine with 1 l/min oxygen throughout the remainder of the procedure. For each animal, a sheath was inserted into the right or left femoral artery, allowing catheter access to the artery and retrograde access to the LV through the aorta. Coronary angiography was performed to locate the left anterior descending (LAD) artery and to confirm patency using the left anterior oblique and/or anterior-posterior views, and the balloon was inflated in the LAD for 90 minutes to simulate MI. Visipaque contrast agent (GE Healthcare) was used to visualize the coronary vasculature using an OEC 9400 c-arm (OEC Medical Systems). The balloon was deflated for 30 minutes to restore blood flow, simulating an angioplasty procedure.

For feasibility testing, 4 or 6 ml of iECM or trilysine with AF568 (n=1 pig per group per volume) was infused using the same PTCA catheter. For dosing analysis, 1 (n=2), 2 (n=3), 4 (n=3), 6 (n=3), 8 (n=3), or 10 (n=2) ml of iECM with AF568 was infused following MI. The pigs from feasibility testing were also used for dosing analysis. For survival procedures, 4 ml of iECM (n=10) or saline (n=10) was infused. The balloon was inflated for 2 minutes, and during that time, up to 2 ml of solution was infused at a rate of approximately 1 ml per minute. The balloon was then deflated for 2 minutes. This cycle of balloon up, infuse 2 ml, and balloon down was repeated 2, 3, 4, or 5 times for a total infusion volume of 4, 6, 8, or 10 ml, respectively. ECG was monitored for signs of arrythmia and collected throughout. LAD patency was confirmed post-MI and post-infusion. For feasibility and dosing analyses, pigs were euthanized 1 hour post-infusion, and heart, lungs, liver, brain, kidney, and spleen were harvested. Hearts were perfused with saline, cut into approximately 1 cm x 1 cm x 2-3 cm rectangular pieces, and embedded in OCT compound for cryosectioning. Two approximately 1 cm x 1 cm x 2-3 cm pieces of each satellite organ (lungs, liver, brain, kidney, spleen) were embedded in OCT and frozen. Samples of each satellite organ were also fixed in 10% formalin for histopathological analysis.

For survival studies, pigs were given banamine (2.2 mg/kg) during recovery. Echocardiography (echo, Siemens Acuson Bonsai ultrasound system), 12-lead electrocardiography (ECG, GE MAC 1200), and Holter monitoring were performed before MI, following infusion, 7±1 days post-op, and 8 weeks ± 2 days post-op (pre-euthanasia), as previously described[3]. Echo and ECG were performed under anesthesia. Holter monitors were attached under anesthesia, pigs were then recovered from anesthesia, and monitors were removed following at least 24 hours, demonstrating at least 12 hours of continuous recording. Holter monitors were reattached if insufficient data (<12h) was collected. Five pigs died during the MI procedure. A standard two-dimensional echocardiogram (2-D Echo) was performed pre-

and post-MI, prior to injection, and prior to euthanasia using a commercially available instrument operating a broad band 1.5-3.5 mHz transducer with digital images stored on hard drive for subsequent playback and analysis. Attempts were made to record images in the standard long axis, short axis, and apical 4 and 2 and 3 chamber views. Depending upon the thoracic configuration of the animal, one or more views that depicted the entire perimeter of the left ventricle were possible. Images were optimized for porcine anatomy with minimal depth and highest frame rate whenever possible, and an acquisition length of 5 cardiac cycles. Once the optimal view was identified, a similar view containing the same landmarks was obtained on each subsequent examination, and single plane LV volumes and ejection fraction were derived using a modified Simpson rule method of discs. Holter recordings were sent to a blinded consultant that prepared full disclosure reports. ECG and Holter reports were evaluated for arrythmias.

**Histological analyses**

All histological and immunohistochemical (IHC) assessment was performed by investigators who were blinded to group assignment. Samples embedded in OCT compound were cryosectioned at 10 and/or 20 μm. Formalin fixed tissue samples were paraffin-embedded and sectioned. Hematoxylin and eosin (H&E)-stained sections were used to identify the infarct regions for fluorescent analyses and infarct size. H&E-stained sections of the pig satellite organs were sent to a histopathologist, who was blinded, and assessed for signs of ischemia and inflammation. Masson’s trichrome stain was used to quantify infarct size and interstitial fibrosis. Five evenly spaced slides spanning the infarct were used for trichrome analyses. Infarcts were manually traced in for infarct size. Slides stained with H&E and trichrome were mounted with Permount (Fisher Chemical) and scanned at 20x using an Aperio Scan Scope CS2 slide scanner (Leica Biosystems). Non-nuclear blue staining was measured using the ‘Positive Pixel Count V9’ algorithm in the ImageScope (Aperio) software for interstitial fibrosis analysis.

IHC was performed using antibodies for the following antigens: α-actinin (αACT, Sigma A7811, 1:700), cleaved caspase 3 ASP175 5A1E (CC3, Cell Signaling 9664, 1:50), and alpha smooth muscle actin (αSMA, Agilent M0851, 1:75), and myeloperoxidase (MPO, Sigma Aldrich N5787, 1:100). Unlabeled antibodies were visualized with secondary antibodies Alexa Fluor 488 or 568 (Life Technologies). Griffonia Simplicifolia Lectin I isolectin B4 (Vector Laboratories, 1:75) was pre-labeled with fluorescein. Nuclei were stained with Hoechst 33342 (Life Technologies). Following IHC, slides were mounted with Fluoromount (Sigma) and imaged at 10X or 20X on the Ariol DM6000 B microscope (Leica) or Carl Zeiss Observer D1.

For neutrophil quantification, MPO was used. Using tissue from 1 day post-infusion, 2 slides covering the infarct were stained. Neutrophils were identified by nuclei colocalization with MPO. Cells were automatically counted using a custom script in Matlab (Mathworks).

For infarct arteriole quantification, arterioles were identified by the following criteria: positive co-staining for isolectin and αSMA for endothelial cells and smooth muscle cells, respectively, a visible lumen, and an average Feret diameter (average of the minimum and maximum Feret diameters) of at least 10 µm. In rats, using tissue from 5 weeks post-infusion, 5 slides evenly spanning the infarct of each heart were stained. In pigs, tissue from 8 weeks post-infusion, 3 slides spanning the infarct of heart were stained. If possible, up to 5 images were captured across the infarct of each slide. Infarcts were identified by nuclei density, and arterioles were manually traced in ImageJ (NIH).

For cardiomyocyte apoptosis quantification in the border zone, slides from 3 days post-infusion were stained for αACT and CC3 to identify cardiomyocytes and apoptotic cells, respectively. The border zone was identified by expanding outside the infarct region 1 mm. The infarct was identified through an absence of αACT stain. Apoptotic cardiomyocytes were identified by positive co-staining for αACT and CC3. As CC3 could be observed on most cells in the border zone, a high threshold for CC3 expression was used. CC3 had to cover at least 50%

of the cell body to be considered apoptotic. Two slides covering the infarct from each heart were used as a result of using the Slice Matrix. Analysis was manually performed in ImageJ (NIH).

For confocal imaging of vessels and iECM, slides from 2, 12, and 24 hours post-infusion in rats and 1 hour post-infusion in pigs were stained with isolectin or αSMA to identify endothelial cells or smooth muscle cells, respectively. Slides were imaged using a Confocal Microscope LSM 780 (Zeiss).

**RNA isolation and NanoString Multiplex Gene Expression Analysis**

Hearts from 1 and 3 days post-infusion were used for gene expression analyses (see Small Animal Surgeries). The LV free wall was isolated and flash frozen for subsequent RNA extraction using the RNEasy Mini Kit (Qiagen). RNA concentrations were measured using a NanoDrop spectrophotometer (Thermo Scientific).

For screening of various pathways of interest, RNA samples were analyzed by NanoString nCounter® MAX Analysis System with nCounter® custom cardiac codeset (Rat) allowing for multiplexed assessment of 380 genes (see Extended Data Table 2 for full gene list). Samples were processed according to manufacturer instructions. In brief, RNA sample concentrations were measured on a Qubit 3.0 Fluorometer with a QubitTM RNA HS Assay kit. 70 µL of hybridization buffer was mixed with Immunology Panel Reporter CodeSet solution, and 8 µL of this master mix was mixed in a separate reaction vessel with 50-100 ng of RNA per tissue sample and RNA-free water up to 13 µL total. 2 µL of Capture ProbeSet was added to each vessel, mixed and placed on a thermocycler at 65ºC for 16-48 hours before being maintained at 4ºC for less than 24 hours. NanoString nCounter Prep Station performed automated fluidic sample processing to purify and immobilize hybridized sample to the cartridge surface. Digital barcode reads were analyzed by NanoString nCounter® Digital Analyzer.

Results were analyzed by manufacturer nSolverTM Analysis Software 4.0 and custom R scripts. Outliers were detected and excluded based on NanoString's methods for outlier detection.

### Statistics

Data generated from dsDNA and sGAG content assays, MRI, qPCR, echocardiography, and histological analyses were compared with a Student's two-way t test. Data generated from hemocompatibility and individual segment wall motion scores were compared with a one-way ANOVA. All data are presented as mean ± SEM. Significance was considered $p < 0.05$. Statistical analysis was performed in Prism (GraphPad).

### Data Availability

**Extended Data Figures**

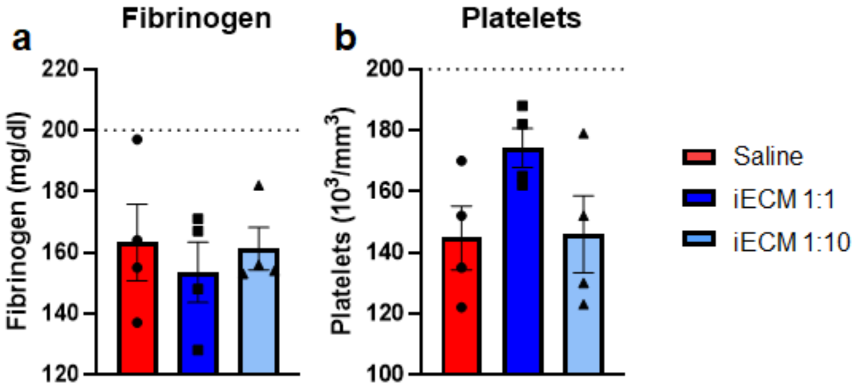


Extended Data Figure 1: iECM does not affect concentration of **a**, fibrinogen and **b**, platelets when mixed with human blood. Standard ranges for each parameter are indicated below dashed lines. N=4 human blood donors. Data are mean ± SEM.

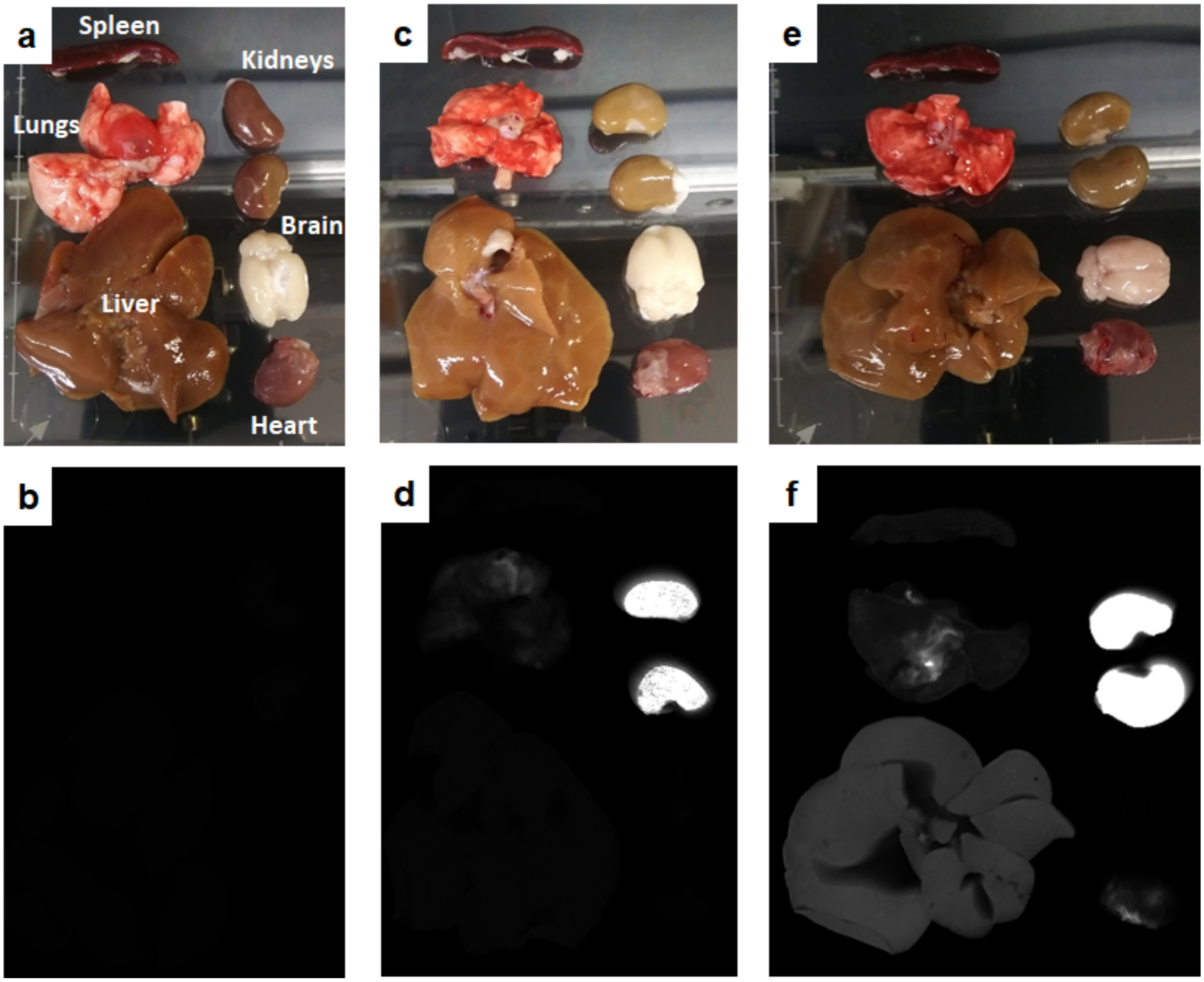


Extended Data Figure 2: iECM biodistribution using VivoTag®-S 750 (VT750) 24 hours following infusion and myocardial infarction procedure. iECM infused rats showed increased signal in the hearts, lungs and liver, with comparable signal in the kidneys, suggesting rapid clearance. Organs labeled in (**a**), the same orientation was used for **c**-**e**. Satellite organs (**a,c,e**), corresponding near-infrared scans, VT750 in white (**b,d,f**). Harvested organs following MI and infusion of saline without VT750 (**a,b**), trylisine with conjugated with VT750 (**c,d**), or iECM conjugated with VT750 (**e,f**).

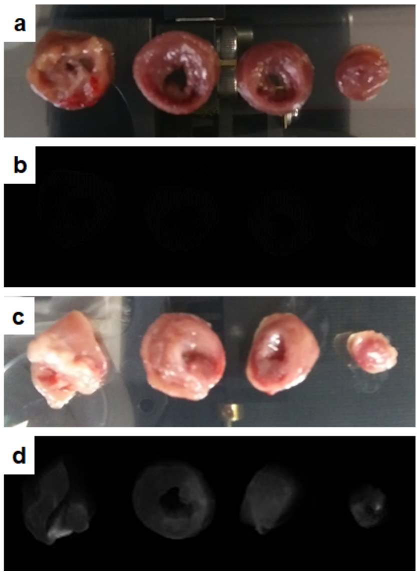


Extended Data Figure 3: Tracking using VivoTag®-S 750 (VT750) 24 hours following MI procedure and infusion of saline without dye (**a**,**b**) or trilysine conjugated with VT750 (**c**,**d**), showing minimal signal as compared to hearts infused with iECM conjugated with VT750 (Fig 2). Images (**a**,**c**) and corresponding near-infrared scans (**b**,**d**) of short axis heart sections. Same scan settings as used in Fig 3.

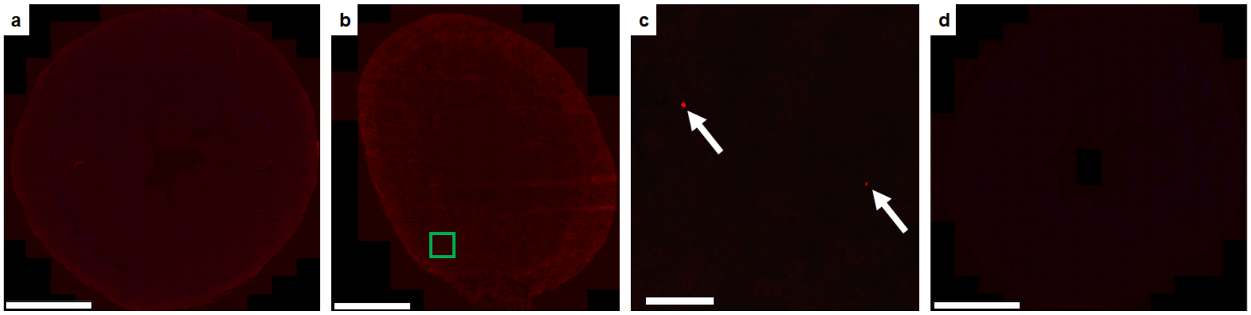


Extended Data Figure 4: Infusible extracellular matrix (iECM) infusions in healthy rats showed no retention in the heart; however, material was observed in the kidneys. **a**, Short axis scan of healthy rat heart following infusion of iECM. Scale bar is 3 mm. **b**, Scan of healthy rat kidney. Scale bar is 2 mm. **c**, Inset of healthy rat kidney. White arrows denoting iECM aggregates. Scale bar is 300 μm. **d**, Short axis scan of chronic MI heart (4 weeks post-MI) following infusion of iECM. Scale bar 3 mm.

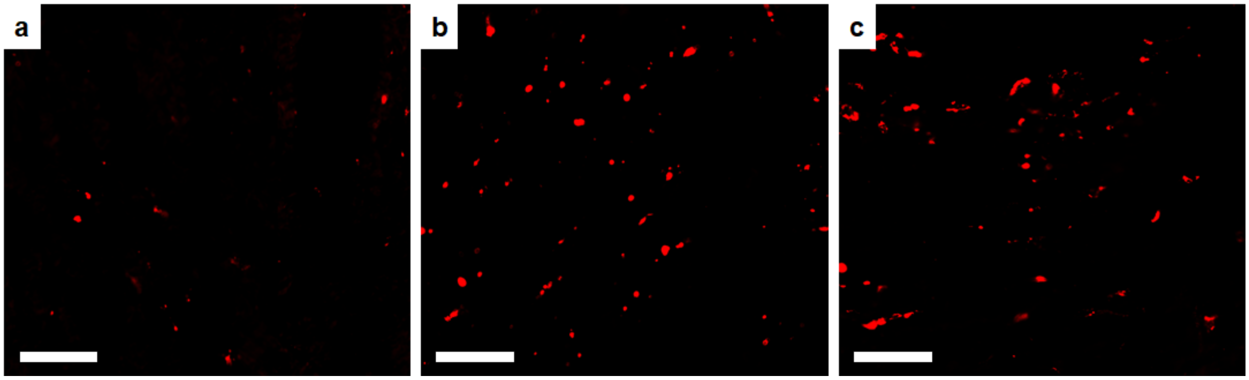


Extended Data Figure 5: *In vivo* dosing of infusible extracellular matrix (iECM) post-MI and subsequent intracoronary infusion showing that high concentrations (10 and 12 mg/ml) infusions have increased signal over lower concentrations (6 mg/ml). Images taken from infarct region. **a**, 6 mg/ml iECM. **b**, 10 mg/ml iECM. **c**, 12 mg/ml iECM. Scale bar is 70 μm.

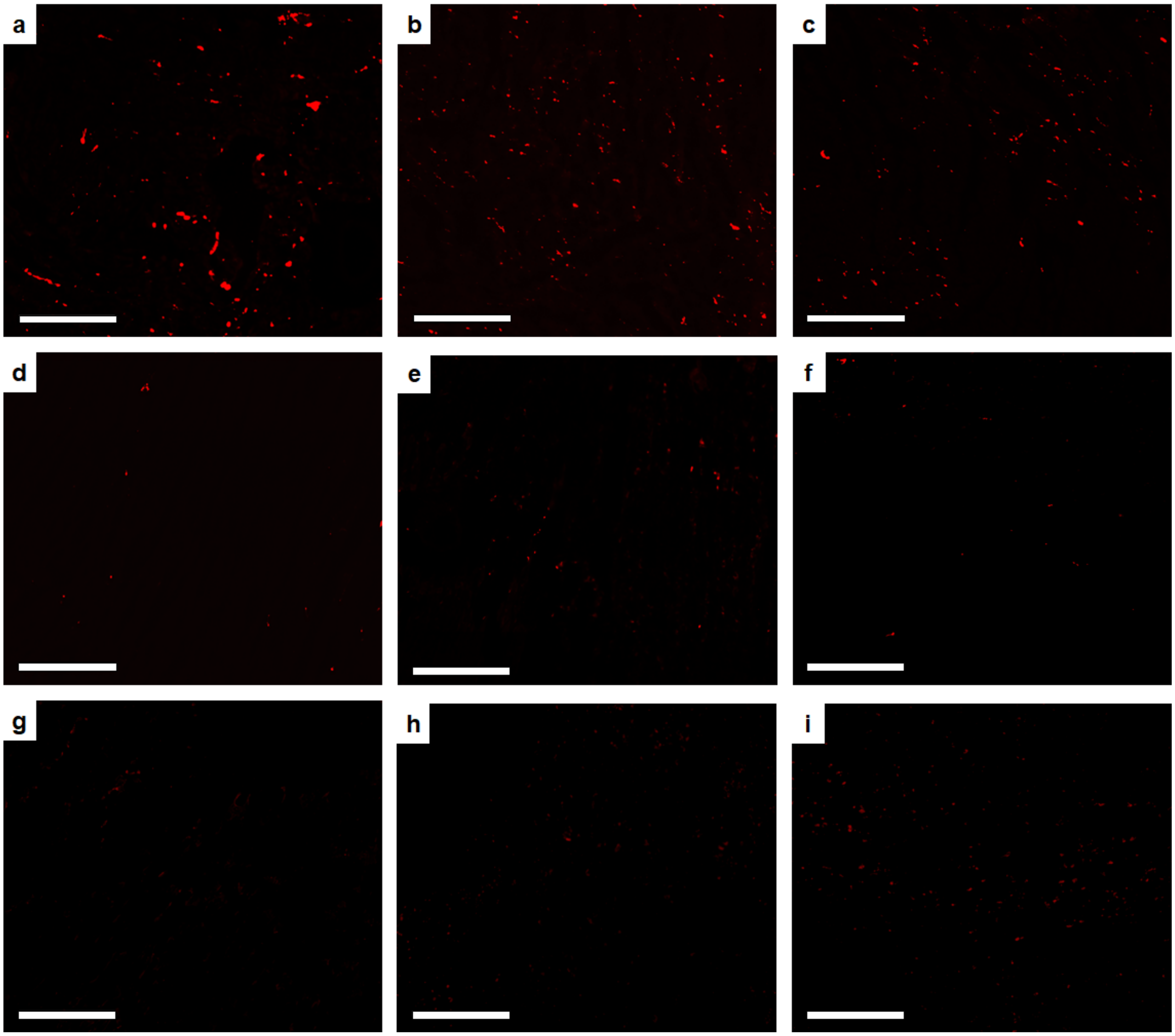


Extended Data Figure 6: *In vivo* degradation of infusible extracellular matrix (iECM) post-MI and subsequent intracoronary infusion showing that iECM degrades in approximately 3 days. Signal could not be differentiated from infarct autofluorescence at 3 days post-infusion. All images taken in the infarct region. **a**, 2 hours post-infusion. **b**, 6 hours post-infusion. **c**, 12 hours post-infusion. **d**, 24 hours post-infusion. **e**, 2 days post-infusion. **f**, 3 days post-infusion. **g**, 4 days post-infusion. **h**, 5 days post-infusion. **i**, 7 days post-infusion. All scale bars are 200 μm.

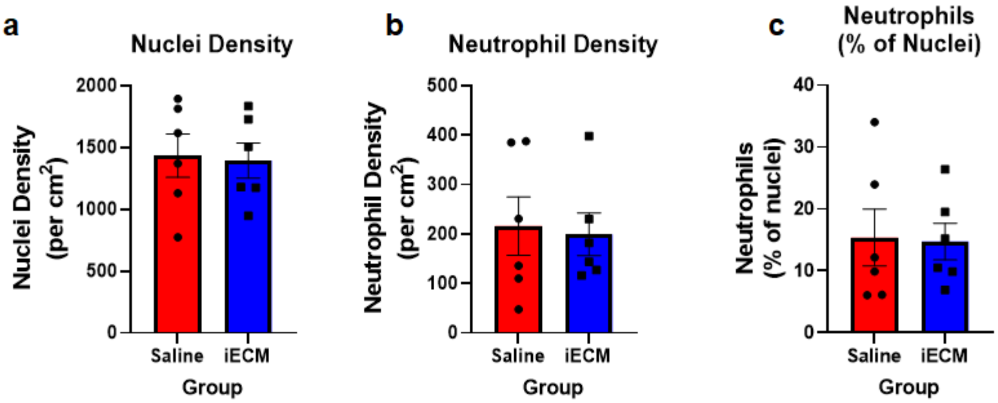


Extended Data Figure 7: Infarct cell density, neutrophil infiltration, and neutrophils as a percentage of total cells were not significantly reduced in response to iECM infusions. Data are mean ± SEM.

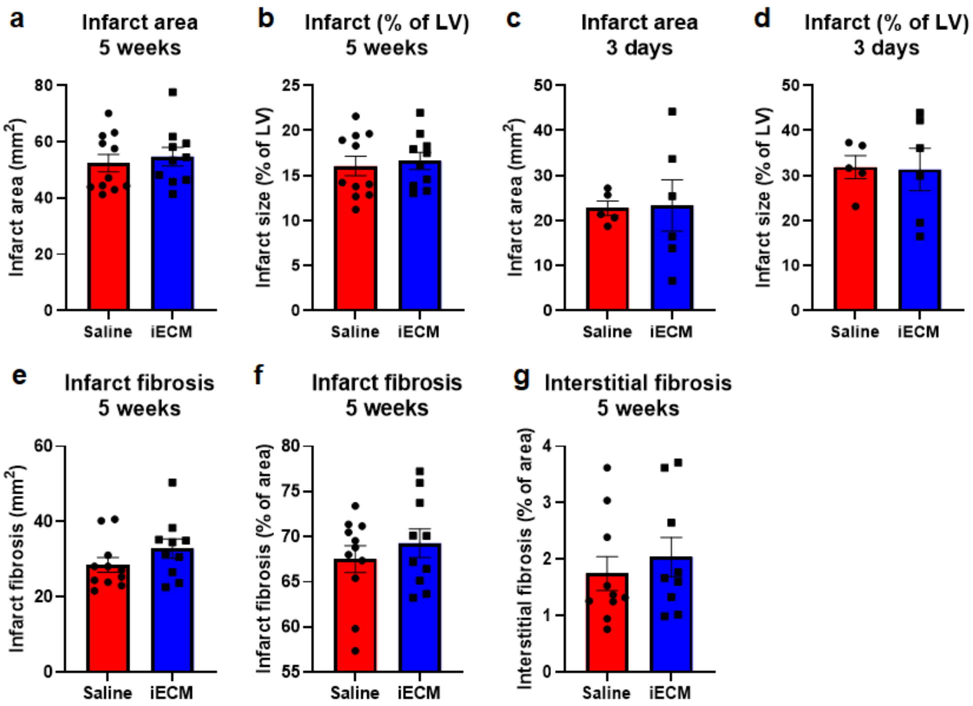


Extended Data Figure 8: Histological measurements post-infusion show no significant differences in infarct size or fibrosis between iECM and saline infused rats. Infarct area at 5 weeks (**a**,**b**) and 3 days (**c**,**d**) post-infusion, reported as area (**a**,**c**) and percentage of the LV (**b**,**d**). Infarct fibrosis reported as area (**e**) and percentage of infarct area (**f**). **g**, Interstitial fibrosis of the remote myocardium reported as a percentage of area. Data are mean ± SEM.

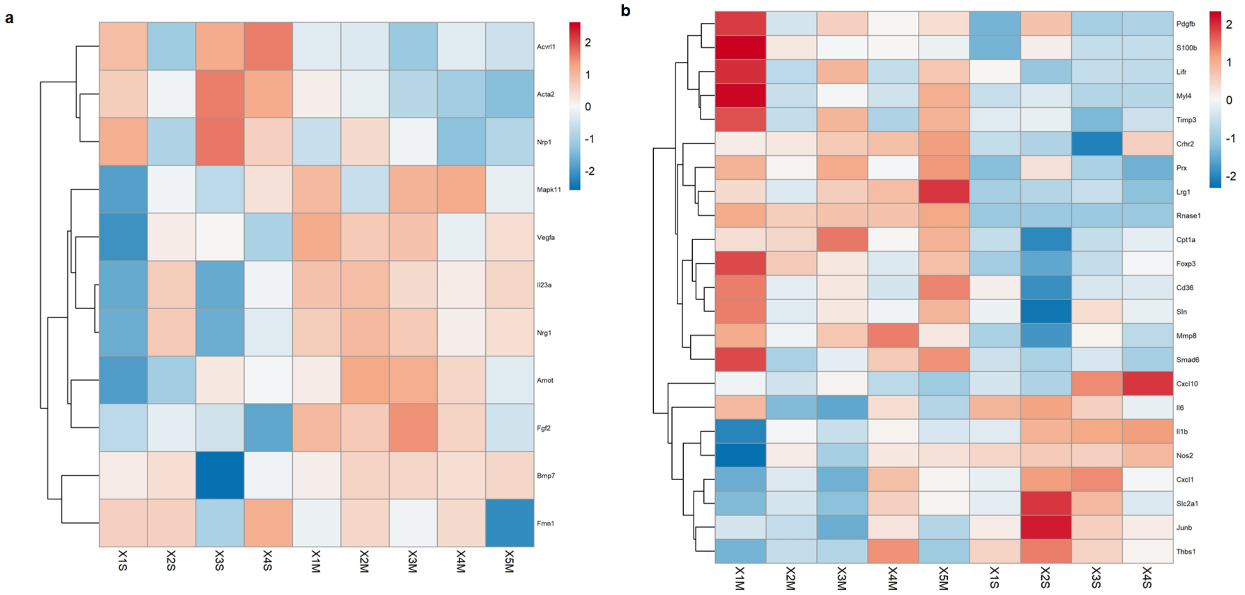


Extended Data Figure 9: Differential gene expression observed at 1 day (**a**) and 3 days (**b**) post-iECM infusion. Heat maps of significantly differentially expressed genes show that gene expression of iECM infused hearts (M) are distinct from saline infused hearts (S).

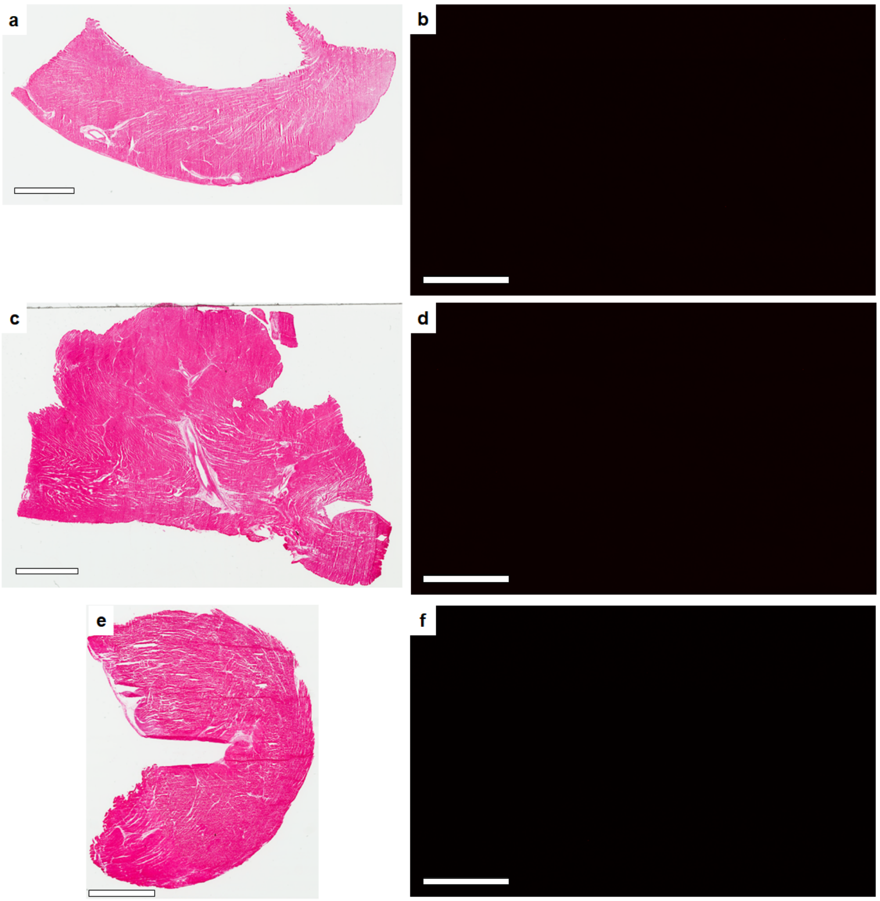


Extended Data Figure 10: Representative H&E (**a**,**c**,**e**) and corresponding representative fluorescent images (**b**,**d**,**f**) of infused pig heart tissue showing no iECM retention in border zone or remote myocardium in addition to no signal in trilysine infused pigs one hour after infusion. **a-b**, Border zone of infarct region of heart infused with iECM conjugated with AF568. **c-d**, Remote myocardium of heart infused with iECM conjugated with AF568. **e-f**, Infarcted region of heart infused with trilysine conjugated with AF568. **a**,**c**,**e**, Scale bar 4 mm. **b**,**d**,**f**, Scale bar 200 μm.

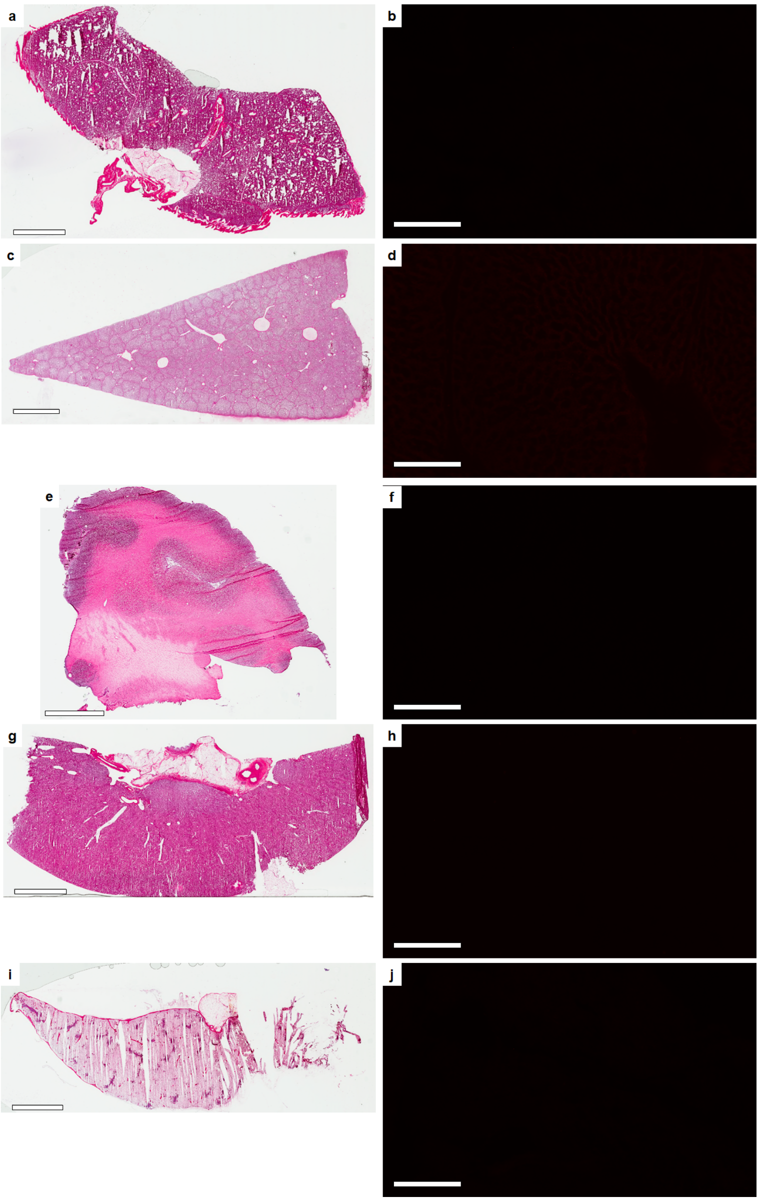

Extended Data Figure 11: Representative H&E (**a**,**c**,**e**,**g**,**i**) and corresponding representative fluorescent images (**b**,**d**,**f**,**h**,**j**) of satellite organs following iECM conjugated with AF568 infusions to the heart showing no observed iECM retention in satellite organs. **a-b**, Lungs. **c-d**, Liver. **e-f**, Brain. **g-h**, Kidney. **i-j**, Spleen. **a**, Scale bar 4 mm. **c**,**e**,**g**, Scale bar 5 mm. **i**, Scale bar 6 mm. **b**,**d**,**f**,**h**,**j**, Scale bars 200 µm.

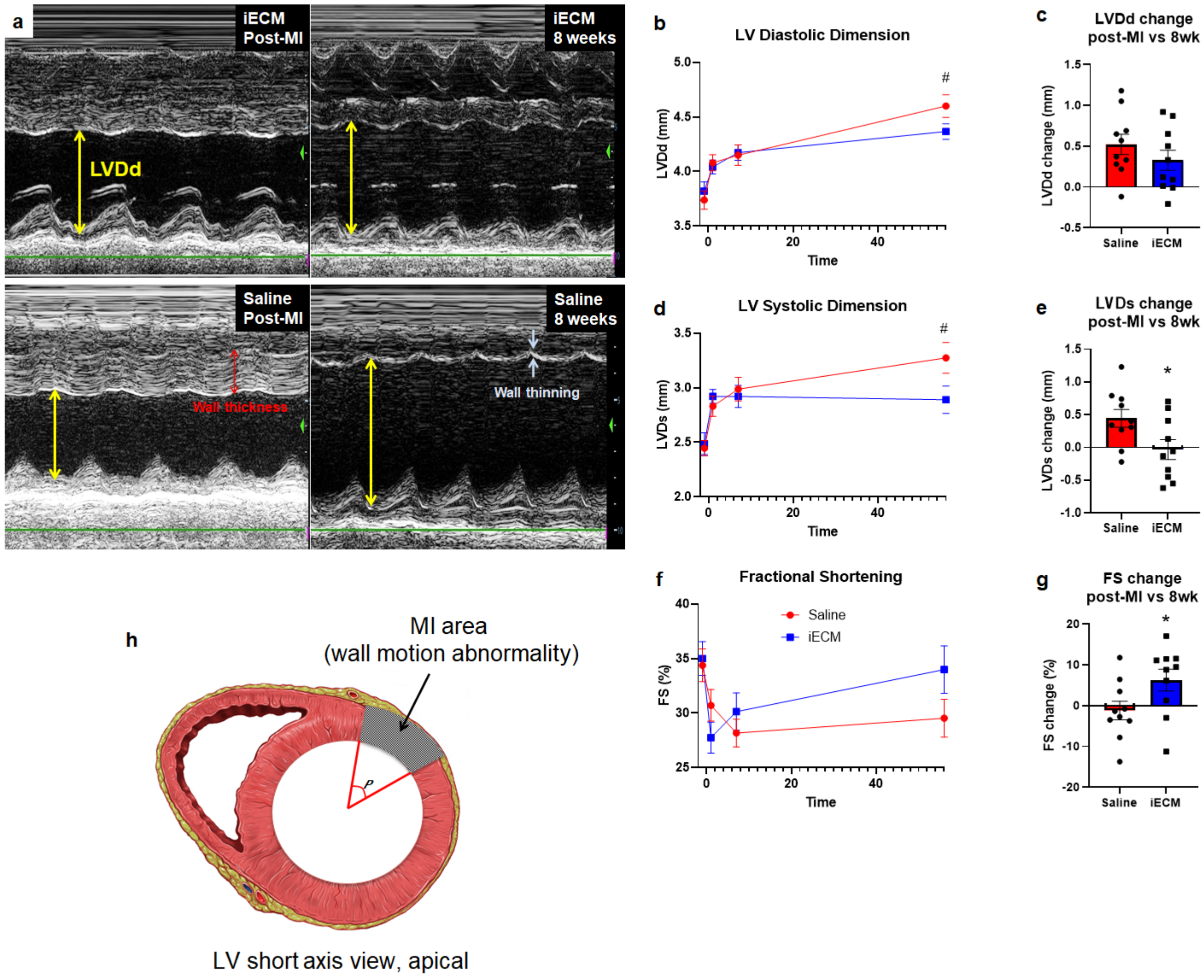


Extended Data Figure 12: Additional echocardiography results showing that iECM infusions mitigate negative LV remodeling in a pig acute MI model. (**a**) Representative M-mode echocardiographic images showing that iECM mitigates negative LV remodeling. Yellow arrows represent LV diastolic dimension, red arrows represent wall thickness, and white arrows represent wall thinning. **b-g**, LV diastolic dimension (LVDd), LV systolic dimension (LVDs), and fractional shortening (FS) over time and changes from post-MI to 8 weeks post-MI. **h**, Diagram demonstrating how infarct angle was measured.

Extended Data Table 1: Design Approach for NanoString nCounter CodeSet

| Pathway | Number of Genes |
|---|---|
| Metabolism | 60 |
| Apoptosis | 23 |
| Neovascularization | 65 |
| Neurogenesis | 24 |
| Immune/Inflammatory Response | 54 |
| Fibrosis | 36 |
| Myogenesis/muscle contraction | 69 |
| Muscle hypertrophy | 44 |
| Housekeeping | 6 |

Extended Data Table 2. Full Gene List for NanoString nCounter CodeSet

| | |
|---|---|
| ABCF1 | NM_001109883.2 |
| Abl1 | NM_001100850.1 |
| Acacb | NM_053922.1 |
| Acly | NM_016987.2 |
| Acox2 | NM_145770.1 |
| Acsm5 | NM_001014162.4 |
| Acta2 | NM_031004.2 |
| Actc1 | NM_019183.1 |
| Actn2 | NM_001170325.1 |
| Actn3 | NM_133424.2 |
| Acvrl1 | NM_022441.2 |
| Adam12 | XM_017590240.1 |
| Adam17 | NM_020306.1 |
| Adgre1 | NM_001007557.1 |
| Adora1 | NM_017155.2 |
| Adrb2 | NM_012492.2 |
| Aggf1 | XM_001060407.6 |
| Agrn | NM_175754.1 |
| Agt | NM_134432.2 |
| Agtr1a | NM_030985.4 |
| Akt1 | NM_033230.1 |
| Akt2 | NM_017093.1 |
| Akt3 | NM_031575.1 |
| Amot | XM_001056974.3 |
| Ang2 | NM_001012359.1 |
| Angptl1 | NM_001109383.1 |
| Angptl3 | NM_001025065.1 |
| Angptl4 | NM_199115.2 |
| Arg1 | NM_017134.2 |
| Artn | NM_053397.1 |
| Atf3 | NM_012912.1 |
| Atf6 | NM_001107196.1 |
| Atp12a | NM_133517.1 |
| Atp1a4 | NM_022848.1 |
| Atp1b1 | NM_013113.2 |
| Atp1b2 | NM_012507.3 |
| Atp1b3 | NM_012913.1 |
| Atp1b4 | NM_053381.1 |
| Atp2a1 | NM_058213.1 |
| Atp2a2 | NM_001110139.2 |
| ATP5F1A | NM_023093.1 |
| Bcl2 | NM_016993.1 |
| Bdh2 | NM_001106473.1 |
| Bmp2 | NM_017178.1 |
| Bmp4 | NM_012827.2 |

| | |
|---|---|
| Bmp7 | NM_001191856.1 |
| Bst1 | NM_030848.1 |
| Btg1 | NM_017258.1 |
| Btk | NM_001007798.1 |
| Cacna1c | NM_012517.2 |
| Cacna1d | NM_017298.1 |
| Cacna2d2 | NM_175592.2 |
| Cacnb2 | NM_053851.1 |
| Cacnb3 | NM_012828.2 |
| Cacnb4 | NM_001105733.1 |
| Cacng1 | NM_019255.1 |
| Cacng4 | NM_080692.1 |
| Cacng5 | NM_080693.1 |
| Calm1 | NM_031969.2 |
| Calm2 | NM_017326.2 |
| Camk2d | NM_012519.2 |
| Camkk1 | NM_031662.1 |
| Camkk2 | NM_031338.1 |
| Casp1 | NM_012762.2 |
| Casp3 | NM_012922.2 |
| Casp4 | NM_053736.2 |
| Casp6 | NM_031775.2 |
| Casp7 | NM_022260.3 |
| Casq2 | NM_017131.2 |
| Cat | NM_012520.1 |
| Ccl1 | NM_001191092.1 |
| Ccl11 | NM_019205.1 |
| Ccl2 | NM_031530.1 |
| Ccl3 | NM_013025.2 |
| Ccl9 | NM_001012357.1 |
| Ccn2 | NM_022266.2 |
| Ccr1 | NM_020542.2 |
| Ccr2 | NM_021866.1 |
| Ccr3 | NM_053958.1 |
| Ccr5 | NM_053960.3 |
| Cd36 | NM_031561.2 |
| Cd40 | NM_134360.1 |
| Cd59 | NM_012925.1 |
| Cd68 | NM_001031638.1 |
| Cdc42 | NM_171994.4 |
| Cflar | NM_057138.2 |
| Chat | NM_001170593.1 |
| Chrm2 | NM_031016.1 |
| Cox4i1 | NM_017202.1 |
| Cox4i2 | NM_053472.1 |
| Cox5b | NM_053586.1 |
| Cox6a1 | NM_012814.1 |
| Cox6a2 | NM_012812.3 |
| Cox6b1 | NM_001145273.1 |

| | |
|---|---|
| Cox6b2 | NM_001039085.1 |
| Cox6c | NM_019360.2 |
| Cox7a2l | NM_001106704.1 |
| Cox7b2 | XM_006221734.3 |
| Cpt1a | NM_031559.2 |
| Cpt1b | NM_013200.1 |
| Creb1 | NM_134443.1 |
| Crhr2 | NM_022714.1 |
| Crtc2 | NM_001033895.1 |
| Cs | NM_130755.1 |
| Csf3 | NM_017104.1 |
| Ctf1 | NM_017129.1 |
| Cxcl1 | NM_030845.1 |
| Cxcl10 | NM_139089.1 |
| Cxcl2 | NM_053647.1 |
| Cxcl3 | NM_138522.1 |
| Cxcl6 | NM_022214.1 |
| Cxcr4 | NM_022205.3 |
| Dffa | NM_053679.2 |
| Dffb | NM_053362.1 |
| Diablo | NM_001008292.1 |
| Dusp5 | NM_133578.1 |
| Dyrk1a | NM_012791.2 |
| Egf | NM_012842.1 |
| Egfl7 | NM_139104.1 |
| Eif2b5 | NM_138866.2 |
| Ep300 | XM_001076610.3 |
| Erbb2 | NM_017003.2 |
| Ereg | NM_021689.1 |
| Esr1 | NM_012689.1 |
| Esrrg | NM_203336.2 |
| Fabp3 | NM_024162.1 |
| Fadd | NM_152937.2 |
| Fas | NM_139194.2 |
| Faslg | NM_012908.1 |
| Fasn | NM_017332.1 |
| Fbp2 | NM_053716.1 |
| Fbxo32 | NM_133521.1 |
| Fcer1a | NM_012724.4 |
| Fgf2 | NM_019305.2 |
| Fh | NM_017005.2 |
| Flt4 | NM_053652.1 |
| Fmn1 | XM_006224644.3 |
| Fos | NM_022197.1 |
| Foxo1 | NM_001191846.2 |
| Foxo3 | NM_001106395.1 |
| Foxp3 | NM_001108250.1 |
| Fst | NM_012561.1 |
| Fxyd2 | NM_017349.2 |

| | |
|---|---|
| G6pd | NM_017006.2 |
| Gadd45a | NM_024127.2 |
| Gata3 | NM_133293.1 |
| Gata4 | NM_144730.1 |
| Gja1 | NM_012567.2 |
| Gls | NM_012569.2 |
| Glud1 | NM_012570.1 |
| Got1 | NM_012571.1 |
| Got2 | NM_013177.2 |
| Gpt | NM_031039.1 |
| Gpt2 | NM_001012057.1 |
| Gpx | NM_030826.2 |
| Gsk3a | NM_017344.1 |
| Gsk3b | NM_032080.1 |
| Gsr | NM_053906.1 |
| Gss | NM_012962.1 |
| GUSB | NM_017015.2 |
| Hey1 | NM_001191845.1 |
| Hey2 | NM_130417.1 |
| Hk2 | NM_012735.1 |
| HPRT1 | NM_012583.2 |
| Hspa5 | NM_013083.1 |
| Idh1 | NM_031510.1 |
| Idh2 | NM_001014161.1 |
| Ifng | NM_138880.2 |
| Igf1 | NM_001082477.2 |
| Il10 | NM_012854.2 |
| Il12a | NM_053390.1 |
| Il13 | NM_053828.1 |
| Il17a | NM_001106897.1 |
| Il1b | NM_031512.1 |
| IL38 | NM_001108571.1 |
| Il1rn | NM_022194.2 |
| Il2 | NM_053836.1 |
| Il23a | NM_130410.2 |
| Il33 | NM_001014166.1 |
| Il36a | NM_001106554.1 |
| Il4 | NM_201270.1 |
| Il5 | NM_021834.1 |
| Il6 | NM_012589.1 |
| Il6st | NM_001008725.3 |
| Irf4 | NM_001106108.1 |
| Irf5 | NM_001106586.1 |
| Itga1 | NM_030994.2 |
| Itga2 | XM_345156.6 |
| Itga3 | XM_003752369.1 |
| Itgav | NM_001106549.1 |
| Itgb1 | NM_017022.2 |
| Itgb3 | NM_153720.1 |

| | |
|---|---|
| Itgb5 | NM_147139.2 |
| Itgb6 | NM_001004263.1 |
| Itgb8 | NM_001108726.1 |
| Jdp2 | NM_053894.1 |
| Junb | NM_021836.2 |
| Kcnd2 | NM_031730.2 |
| Kdr | NM_013062.1 |
| Kit | NM_022264.1 |
| Klf10 | NM_031135.2 |
| Kmt2a | XM_006226455.1 |
| LDHA | NM_017025.1 |
| Lep | NM_013076.3 |
| Letm1 | NM_001005884.1 |
| Lifr | NM_031048.1 |
| Lipe | NM_012859.1 |
| Lox | NM_017061.2 |
| Lrg1 | NM_001009717.1 |
| Map2k3 | XM_001077724.1 |
| Mapk1 | NM_053842.1 |
| Mapk11 | NM_001109532.2 |
| Mapk12 | NM_021746.1 |
| Mapk14 | NM_031020.2 |
| Mapk8 | XM_001056513.1 |
| Mapkapk2 | NM_178102.2 |
| Mcl1 | NM_021846.2 |
| Mcu | NM_001106398.1 |
| Mdh1 | NM_033235.1 |
| Mdk | NM_030859.2 |
| Mef2c | XM_006223957.2 |
| Mef2d | NM_030860.2 |
| Mmp1 | NM_001134530.1 |
| Mmp10 | NM_133514.1 |
| Mmp12 | NM_053963.1 |
| Mmp13 | NM_133530.1 |
| Mmp14 | NM_031056.1 |
| Mmp19 | NM_001107159.1 |
| Mmp2 | NM_031054.2 |
| Mmp25 | XM_001055465.2 |
| Mmp28 | NM_001079888.1 |
| Mmp3 | NM_133523.1 |
| Mmp8 | NM_022221.1 |
| Mmp9 | NM_031055.1 |
| Mstn | NM_019151.1 |
| Myd88 | NM_198130.1 |
| Myef2 | NM_001013205.2 |
| Myh6 | NM_017239.2 |
| Myh7 | NM_017240.1 |
| Myh8 | NM_001100485.1 |
| Myl2 | NM_001035252.2 |

| | |
|---|---|
| Myl3 | NM_012606.2 |
| Myl4 | NM_001109495.1 |
| Myocd | NM_182667.2 |
| Nf1 | NM_012609.1 |
| Nfatc4 | NM_001107264.1 |
| Nfkb1 | XM_342346.3 |
| Nkg7 | NM_133540.1 |
| Nkx2-5 | NM_053651.1 |
| Nog | NM_012990.1 |
| Nol3 | NM_053516.2 |
| Nos2 | NM_012611.2 |
| Nos3 | NM_021838.2 |
| Notch2 | NM_024358.1 |
| Nr4a1 | NM_024388.2 |
| Nrg1 | NM_001271120.1 |
| Nrp1 | NM_145098.2 |
| Nrp2 | NM_030869.3 |
| Nupr1 | NM_053611.1 |
| Pax2 | NM_001106361.1 |
| Pax3 | NM_053710.1 |
| Pc | NM_012744.2 |
| Pdgfa | NM_012801.1 |
| Pdgfb | NM_031524.1 |
| Pdgfc | NM_031317.1 |
| Pdha1 | NM_001004072.2 |
| Pdk1 | NM_053826.2 |
| Pdk2 | NM_030872.1 |
| Pdk3 | NM_001106581.1 |
| Pdk4 | NM_053551.1 |
| Pecam1 | NM_031591.1 |
| Pecr | NM_133299.1 |
| Pf4 | NM_001007729.1 |
| Pgf | NM_053595.2 |
| Pik3ca | XM_001059296.1 |
| Pik3r1 | NM_013005.1 |
| Pkm | NM_053297.2 |
| Pla2g12a | NM_001108565.1 |
| Pla2g5 | NM_017174.1 |
| POLR1B | NM_031773.1 |
| Ppargc1a | NM_031347.1 |
| Ppargc1b | NM_176075.2 |
| Ppbp | NM_153721.1 |
| Ppp3ca | NM_017041.1 |
| Prkaa1 | NM_019142.1 |
| Prkaa2 | NM_023991.1 |
| Prkca | XM_343975.3 |
| Prkcb | NM_001172305.1 |
| Prkce | NM_017171.1 |
| Prkcg | NM_012628.1 |

| | |
|---|---|
| Prl7d1 | NM_053364.1 |
| Prx | NM_023976.2 |
| Ptgs1 | NM_017043.3 |
| Ptk2 | NM_013081.2 |
| Ptn | NM_017066.2 |
| Pxn | NM_001012147.1 |
| Rac1 | NM_134366.1 |
| Rac2 | NM_001008384.1 |
| Raf1 | NM_012639.2 |
| Rheb | NM_013216.1 |
| Rhoa | NM_057132.3 |
| Rnase1 | NM_001029904.1 |
| RPLP0 | NM_022402.2 |
| Runx1 | NM_017325.1 |
| Ryr2 | NM_032078.2 |
| S100b | NM_013191.1 |
| S1pr1 | XM_008761459.1 |
| Sds | NM_053962.3 |
| Serpine1 | NM_012620.1 |
| Shh | NM_017221.1 |
| Ski | XM_017593893.1 |
| Slc2a1 | NM_138827.1 |
| Slc2a4 | NM_012751.1 |
| Slc8a1 | NM_001270772.1 |
| Slc8a2 | NM_078619.1 |
| Slc8a3 | NM_078620.1 |
| Slc9a1 | NM_012652.1 |
| Slit2 | NM_022632.2 |
| Sln | NM_001013247.1 |
| Smad2 | NM_001277450.1 |
| Smad3 | NM_013095.2 |
| Smad4 | NM_019275.2 |
| Smad6 | NM_001109002.2 |
| Smad7 | NM_030858.1 |
| Smo | NM_012807.1 |
| Sod1 | NM_017050.1 |
| Sphk1 | NM_001270811.1 |
| Stat3 | NM_012747.2 |
| Stk11 | NM_001108069.1 |
| Tbx20 | NM_001108132.1 |
| Tbx21 | NM_001107043.1 |
| Tbx5 | NM_001009964.1 |
| Tcap | NM_001271277.1 |
| Tek | NM_001105737.1 |
| Tgfa | NM_012671.2 |
| Tgfb1 | NM_021578.2 |
| Tgfb2 | NM_031131.1 |
| Tgfb3 | NM_013174.2 |
| Tgfbr1 | NM_012775.2 |

| | |
|---|---|
| Tgfbr2 | NM_031132.3 |
| Thbs1 | NM_001013062.1 |
| Tie1 | NM_053545.1 |
| Timp1 | NM_053819.1 |
| Timp2 | NM_021989.2 |
| Timp3 | NM_012886.2 |
| Timp4 | NM_001109393.1 |
| Tlr2 | NM_198769.2 |
| Tlr4 | NM_019178.1 |
| Tnfrsf1a | NM_013091.1 |
| Tnnc1 | NM_001034105.1 |
| Tnni3 | NM_017144.1 |
| Tnnt2 | NM_012676.1 |
| Tp53 | NM_030989.3 |
| Tpm1 | NM_001034068.1 |
| Tpm2 | NM_001024345.1 |
| Tpm3 | NM_173111.1 |
| Tpm4 | NM_012678.2 |
| Tpt1 | NM_053867.1 |
| Traf2 | NM_001107815.2 |
| Traf3 | NM_001108724.1 |
| Trx1 | NM_053800.3 |
| Txnrd1 | NM_031614.2 |
| Tymp | NM_001012122.1 |
| Ucn3 | NM_001080208.1 |
| Uqcr10 | NM_001170465.1 |
| Uqcrb | NM_001127553.2 |
| Uqcrc1 | NM_001004250.2 |
| Uqcrc2 | NM_001006970.1 |
| Uqcrfs1 | NM_001008888.1 |
| Uqcrh | NM_001009480.1 |
| Uqcrq | NM_001025134.1 |
| Vcam1 | NM_012889.1 |
| Vegfa | NM_031836.2 |
| Vegfb | NM_053549.1 |
| Vegfc | NM_053653.1 |
| Wars2 | NM_001168641.1 |
| Wt1 | NM_031534.2 |
| Xiap | NM_022231.2 |
| Yap1 | NM_001034002.2 |

Extended Data Table 3: Top 30 Gene Ontology pathways and the associated differentially expressed genes 1 day post-myocardial infarction and iECM infusion

| ID | Description | qvalue | geneID |
|---|---|---|---|
| GO:0043535 | regulation of blood vessel endothelial cell migration | 3.04E-07 | Acvrl1/Amot/Fgf2/Nrp1/Vegfa |
| GO:0043534 | blood vessel endothelial cell migration | 6.86E-07 | Acvrl1/Amot/Fgf2/Nrp1/Vegfa |
| GO:0090130 | tissue migration | 8.95E-07 | Acta2/Acvrl1/Amot/Fgf2/Nrp1/Vegfa |
| GO:1901342 | regulation of vasculature development | 1.11E-06 | Acvrl1/Amot/Bmp7/Fgf2/Nrp1/Vegfa |
| GO:0010594 | regulation of endothelial cell migration | 1.11E-06 | Acvrl1/Amot/Fgf2/Nrp1/Vegfa |
| GO:0043536 | positive regulation of blood vessel endothelial cell migration | 1.11E-06 | Amot/Fgf2/Nrp1/Vegfa |
| GO:0051893 | regulation of focal adhesion assembly | 1.11E-06 | Acvrl1/Fmn1/Nrp1/Vegfa |
| GO:0090109 | regulation of cell-substrate junction assembly | 1.11E-06 | Acvrl1/Fmn1/Nrp1/Vegfa |
| GO:0150116 | regulation of cell-substrate junction organization | 1.11E-06 | Acvrl1/Fmn1/Nrp1/Vegfa |
| GO:0045785 | positive regulation of cell adhesion | 1.86E-06 | Bmp7/Fmn1/Il23a/Nrg1/Nrp1/Vegfa |
| GO:0043542 | endothelial cell migration | 1.98E-06 | Acvrl1/Amot/Fgf2/Nrp1/Vegfa |
| GO:0048041 | focal adhesion assembly | 2.52E-06 | Acvrl1/Fmn1/Nrp1/Vegfa |
| GO:0010632 | regulation of epithelial cell migration | 2.99E-06 | Acvrl1/Amot/Fgf2/Nrp1/Vegfa |
| GO:0007044 | cell-substrate junction assembly | 3.36E-06 | Acvrl1/Fmn1/Nrp1/Vegfa |
| GO:0150115 | cell-substrate junction organization | 3.36E-06 | Acvrl1/Fmn1/Nrp1/Vegfa |
| GO:0048738 | cardiac muscle tissue development | 4.08E-06 | Bmp7/Fgf2/Mapk11/Nrg1/Vegfa |
| GO:0010595 | positive regulation of endothelial cell migration | 4.39E-06 | Amot/Fgf2/Nrp1/Vegfa |
| GO:0003007 | heart morphogenesis | 5.40E-06 | Acvrl1/Bmp7/Nrg1/Nrp1/Vegfa |
| GO:0010631 | epithelial cell migration | 5.52E-06 | Acvrl1/Amot/Fgf2/Nrp1/Vegfa |
| GO:0090132 | epithelium migration | 5.52E-06 | Acvrl1/Amot/Fgf2/Nrp1/Vegfa |
| GO:0001952 | regulation of cell-matrix adhesion | 6.34E-06 | Acvrl1/Fmn1/Nrp1/Vegfa |
| GO:0045765 | regulation of angiogenesis | 6.36E-06 | Acvrl1/Amot/Fgf2/Nrp1/Vegfa |
| GO:0010634 | positive regulation of epithelial cell migration | 1.22E-05 | Amot/Fgf2/Nrp1/Vegfa |
| GO:0050921 | positive regulation of chemotaxis | 1.29E-05 | Fgf2/Il23a/Nrp1/Vegfa |
| GO:0034329 | cell junction assembly | 1.66E-05 | Acvrl1/Fmn1/Nrg1/Nrp1/Vegfa |
| GO:0048638 | regulation of developmental growth | 1.98E-05 | Fgf2/Mapk11/Nrg1/Nrp1/Vegfa |
| GO:0032103 | positive regulation of response to external stimulus | 2.30E-05 | Fgf2/Il23a/Nrg1/Nrp1/Vegfa |
| GO:0045766 | positive regulation of angiogenesis | 2.49E-05 | Acvrl1/Fgf2/Nrp1/Vegfa |
| GO:0048754 | branching morphogenesis of an epithelial tube | 2.49E-05 | Bmp7/Fgf2/Nrp1/Vegfa |
| GO:0042060 | wound healing | 2.75E-05 | Acta2/Acvrl1/Fgf2/Nrg1/Vegfa |

Extended Data Table 4: Top 30 Gene Ontology pathways and the associated differentially expressed genes 3 days post-myocardial infarction and iECM infusion

| ID | Description | qvalue | geneID |
|---|---|---|---|
| GO:2000379 | positive regulation of reactive oxygen species metabolic process | 2.66E-08 | Cd36/Cxcl1/Il1b/Il6/Mmp8/Pdgfb/Thbs1 |
| GO:0032635 | interleukin-6 production | 3.16E-07 | Cd36/Crhr2/Foxp3/Il1b/Il6/Mmp8/Nos2 |
| GO:0072593 | reactive oxygen species metabolic process | 3.39E-07 | Cd36/Cxcl1/Il1b/Il6/Mmp8/Nos2/Pdgfb/Thbs1 |
| GO:2000377 | regulation of reactive oxygen species metabolic process | 7.99E-07 | Cd36/Cxcl1/Il1b/Il6/Mmp8/Pdgfb/Thbs1 |
| GO:0032675 | regulation of interleukin-6 production | 5.12E-06 | Cd36/Crhr2/Foxp3/Il1b/Il6/Mmp8 |
| GO:0006809 | nitric oxide biosynthetic process | 5.62E-06 | Cd36/Il1b/Il6/Mmp8/Nos2 |
| GO:0046209 | nitric oxide metabolic process | 6.05E-06 | Cd36/Il1b/Il6/Mmp8/Nos2 |
| GO:2001057 | reactive nitrogen species metabolic process | 6.83E-06 | Cd36/Il1b/Il6/Mmp8/Nos2 |
| GO:1901342 | regulation of vasculature development | 8.35E-06 | Cd36/Crhr2/Cxcl10/Il1b/Lrg1/Pdgfb/Thbs1 |
| GO:0032755 | positive regulation of interleukin-6 production | 1.10E-05 | Cd36/Crhr2/Il1b/Il6/Mmp8 |
| GO:0045429 | positive regulation of nitric oxide biosynthetic process | 1.60E-05 | Cd36/Il1b/Il6/Mmp8 |
| GO:1904407 | positive regulation of nitric oxide metabolic process | 1.79E-05 | Cd36/Il1b/Il6/Mmp8 |
| GO:1901343 | negative regulation of vasculature development | 1.94E-05 | Cd36/Crhr2/Cxcl10/Pdgfb/Thbs1 |
| GO:1903409 | reactive oxygen species biosynthetic process | 1.98E-05 | Cd36/Il1b/Il6/Mmp8/Nos2 |
| GO:0050663 | cytokine secretion | 2.16E-05 | Cd36/Foxp3/Il1b/Il6/Mmp8/Nos2 |
| GO:0042060 | wound healing | 3.02E-05 | Cd36/Il1b/Il6/Nos2/Pdgfb/Thbs1/Timp3 |
| GO:1903428 | positive regulation of reactive oxygen species biosynthetic process | 3.30E-05 | Cd36/Il1b/Il6/Mmp8 |
| GO:0001819 | positive regulation of cytokine production | 3.57E-05 | Cd36/Crhr2/Foxp3/Il1b/Il6/Mmp8/Thbs1 |
| GO:0045765 | regulation of angiogenesis | 4.71E-05 | Cd36/Crhr2/Cxcl10/Il1b/Lrg1/Thbs1 |
| GO:0045428 | regulation of nitric oxide biosynthetic process | 4.83E-05 | Cd36/Il1b/Il6/Mmp8 |
| GO:0070372 | regulation of ERK1 and ERK2 cascade | 6.04E-05 | Cd36/Crhr2/Il1b/Il6/Pdgfb/Timp3 |
| GO:0050867 | positive regulation of cell activation | 7.49E-05 | Foxp3/Il1b/Il6/Mmp8/Pdgfb/Thbs1 |
| GO:0070371 | ERK1 and ERK2 cascade | 7.78E-05 | Cd36/Crhr2/Il1b/Il6/Pdgfb/Timp3 |
| GO:0002718 | regulation of cytokine production involved in immune response | 8.44E-05 | Cd36/Foxp3/Il1b/Il6 |
| GO:0015908 | fatty acid transport | 1.00E-04 | Cd36/Il1b/Nos2/Thbs1 |
| GO:1904950 | negative regulation of establishment of protein localization | 1.00E-04 | Cd36/Crhr2/Foxp3/Il1b/Il6 |
| GO:0019221 | cytokine-mediated signaling pathway | 1.00E-04 | Cxcl1/Cxcl10/Il1b/Il6/Lifr/Pdgfb |
| GO:0019935 | cyclic-nucleotide-mediated signaling | 1.18E-04 | Cd36/Crhr2/Cxcl10/Nos2/Thbs1 |

| | | | |
|---|---|---|---|
| GO:0042035 | regulation of cytokine biosynthetic process | 1.25E-04 | Foxp3/Il1b/Il6/Thbs1 |
| GO:0050707 | regulation of cytokine secretion | 1.31E-04 | Cd36/Foxp3/Il1b/Il6/Mmp8 |

Extended Data Table 5: Histopathological assessment of pig satellite organs. Normal (N), Abnormal (Ab), Autolysis (Aut).

| | Infusion and volume | | | |
|---|---|---|---|---|
| Tissue | iECM 4 ml | iECM 6 ml | Trilysine 4 ml | Trilysine 6 ml |
| Lungs | N | N | N | N |
| Liver | N | N | N | N |
| Brain | N | N | N | N |
| Kidney | N | N | N | N |
| Spleen | N | N | N | N |
